# The Innate Immune Toll-Like Receptor-2 modulates the Depressogenic and Anorexiolytic Neuroinflammatory Response in Obstructive Sleep Apnoea

**DOI:** 10.1101/2019.12.24.888206

**Authors:** Dora Polsek, Diana Cash, Mattia Veronese, Katarina Ilic, Tobias C. Wood, Milan Milosevic, Svjetlana Kalanj-Bognar, Mary J. Morrell, Steve C.R. Williams, Srecko Gajovic, Guy D. Leschziner, Dinko Mitrecic, Ivana Rosenzweig

## Abstract

**Background:** The neurological mechanisms of the disease process of obstructive sleep apnea, the second most frequent sleep disorder, remain unclear whilst its links with several major neuropsychiatric disorders, such as depression, anxiety and even Alzheimer’s disorder, are increasingly recognised. A radical theory, that inflammation in the brain may underlie certain phenotypes of many of these disorders, has been proposed, and the microglial TLR2 system may serve as an important crossroad at the borderlands of several pathogenesis. This study undertook to investigate whether a neuroinflammatory response occurs under conditions of OSA, and whether it might be related to a modulated response due to TLR2 functionality in an established rodent model of OSA.

**Methods:** The effects of three weeks’ exposure to chronic intermittent hypoxia were monitored in mice with or without functional TLR2 (C57BL/6-Tyrc-Brd-Tg(Tlr2-luc/gfp)Kri/Gaj; TLR2^−/−^, C57BL/6-Tlr2tm1Kir), that were investigated by multimodal *in vivo* and *ex vivo* imaging, combining magnetic resonance and bioluminescence imaging and a variety of functional tests.

**Results:** An acute neuroinflammatory response was demonstrated following the three days in the basal forebrain of mice, and more chronically in other parts of the frontal cortex. Adaptive changes in specific neurocircuitry were demonstrated, with significant links to agitated (mal)adaptive behaviour under episodes of stress, and an increased ability to gain weight.

**Conclusions:** Our results suggest that microglial activation and an innate immune response might be the missing link underlying the pathogenesis of well known structural, psychologic and metabolic changes experienced by some patients with OSA.

## Introduction

Obstructive sleep apnoea (OSA) is a major clinical problem due to its high prevalence and serious complications(1, 2), including its links to anxiety disorders, depression(2, 3) and Alzheimer’s disease (AD)(4–7). OSA results from a mix of genetic, environmental, and lifestyle factors, with obesity and aging being the key risk factors. In patients with OSA, the upper airway narrows or collapses repeatedly during sleep, causing obstructive apnoeic events associated with intermittent hypoxia, recurrent arousals and increase in respiratory effort, leading to secondary sympathetic activation, oxidative stress and systemic inflammation.(8) To date, the neurological mechanisms of the disease process of OSA remain unclear. Their deciphering is of paramount importance, as it might aid the development of an effective neuroprotective therapeutic approach in AD, with which OSA appears to share a complex bidirectional link(4, 9). Whilst several studies suggest that glial activation and associated inflammation play an important role in the pathogenesis of AD, depression(10, 11) and several other psychiatric disorders(12), the occurrence of any neuroinflammatory process is yet to be demonstrated in OSA.

Toll-like receptors (TLRs) serve as important links between innate and adaptive immunity of the brain, and most recently the microglial TLR2 system has been demonstrated to play an important role in AD pathogenesis(13). Similarly, several recent pivotal studies suggested abnormalities in TLRs, including the TLR2 system, might be playing an important role in the pathophysiology of depression and suicidal behaviour(14), stress-induced neuroinflammation(15), elevated anxiety and social avoidance(16).

To investigate whether a neuroinflammatory response occurs under conditions of OSA, and to elucidate to what extent the combination of beneficial and harmful inflammatory events might be related to a modulated response due to TLR2 functionality, this study monitored the effects of three weeks’ exposure to chronic intermittent hypoxia (IH) in an established rodent model of OSA(17),(18). This mouse model has been shown to verifiably mimic the electroencephalographic arousals and significant hypoxaemia experienced by patients with OSA. Here its consequences were studied through time by multimodal *in vivo* and *ex vivo* imaging, combining magnetic resonance imaging (MRI) and bioluminescence imaging (BLI). The imaging was complemented by functional tests (weight monitoring, Y-maze, open field and tail suspension behavioural tests), imaging *versus* mRNA expression analysis and *ex vivo* analyses of variety of cellular markers (Supplement).

## Methods

Two mouse lines were used, C57BL/6-Tyr^c-Brd^-Tg(Tlr2-luc/gfp)^Kri^/Gaj and C57BL/6-*Tlr2^tm1Kir^*, (further in the text TLR2 and TLR2^−/−^ respectively), both previously described by our collaborators(19) (see Supplemental Material and Methods). Four experimental groups were compared: group 1 (TLR2IH), mice with functional TLR2 system that were exposed to three weeks of chronic IH protocol(18, 20); group 2 (TLR2CTRL), mice with functional TLR2-system that were handled under control (CTRL) conditions; group 3 (TLR2^−/−^ IH), TLR2 knock out mice exposed to three weeks of chronic IH; and group 4 (TLR2^−/−^ CTRL) control TLR2 knock out mice. In BLI experiments we compared 2 groups (TLRIH and TLR2CTRL) of mice, and all other investigations were done between 4 groups (TLR2IH, TLR2CTRL, TLR2^−/−^IH and TLR2^−/−^CTRL) (Supplemental Figure S1 depicts research protocol). Detailed methodological description of mice lines, all experimental and study procedures, including the undertaken statistical and MR analyses, is available in the Supplement.

## Results

### An acute two-site TLR2 response in mouse model of OSA

Previous studies have shown that TLR2 regulates the hypoxic/ischaemic brain damage caused by stroke(4, 19, 21). To establish whether IH, that results from nocturnal apnoeic and hypopnoeic episodes in patients with OSA, provokes a similar inflammatory response in brain, we used an established mouse model of OSA(18, 20) and transgenic mice that bore the dual reporter system luciferase/green fluorescent protein under transcriptional control of the murine TLR2-promoter. TLR2 induction/microglial activation and its spatial and temporal dynamics were then investigated longitudinally in real time using BLI, as previously described(19). The signals were analysed over a three-week period following experimental (IH) and control (CTRL) protocol (Figure 1A-D).

**Figure 1.**
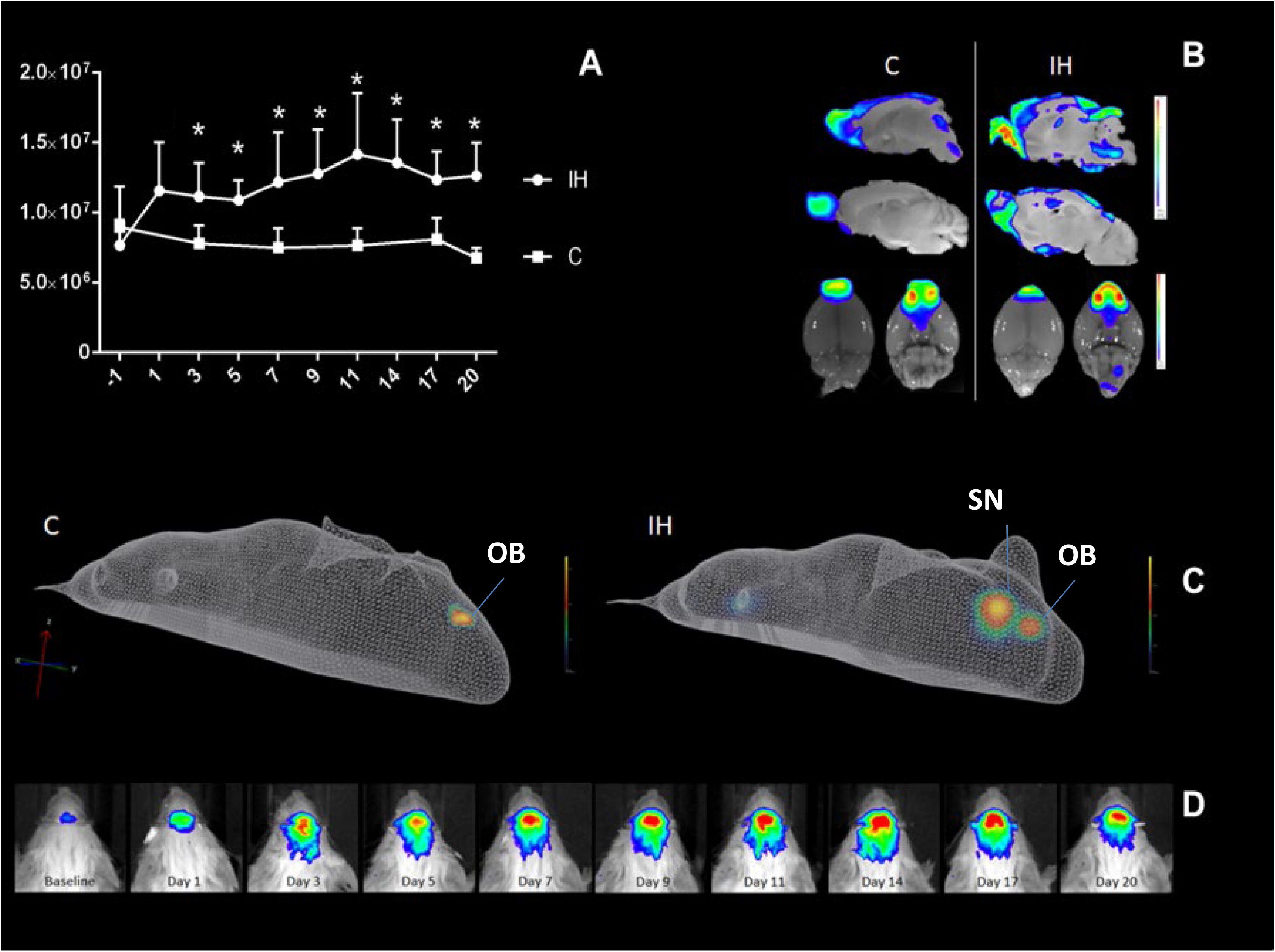
Neuroinflammatory response in a rodent model of obstructive sleep apnoea. Real-time imaging of TLR2 response following intermittent hypoxia reveals a long-term induction of inflammatory signals. (**A–D**) *In vivo* imaging of TLR2 induction after intermittent hypoxia shows activation of the TLR2 promoter up to three weeks (**A, D**). **A:** Plot of the data obtained by measuring the luciferase activity (in photon per second, p/s) in control (C; n=6) and intermittent hypoxia group (IH; n=7) is depicted. Quantification of bioluminescense (BLI) signal in exposed and control groups (IH and C respectively) shows statistically significant increase of neuroinflammatory response after one day of IH that remains elevated throughout the protocol. *X axis* shows BLI signal in photons/sec/cm2/sr, *Y axis* depicts day of IH protocol. The solid white line shows the TLR2 induction: note the strong induction of the promoter within 72 hours of protocol, with a sub/chronic induction and continued significant expression at twenty days (mean ± SD; Wilcoxon signed ranks test, *P* < .05 *compared with baseline values). A representative image of *ex vivo* TLR2 signal following one day of protocol is shown (**B**). The signal is localized to the olfactory bulbs and anterior olfactory nucleus (OB) of both treatment groups at baseline, but intensifies and spreads posteriorly already after only one day of exposure to intermittent hypoxia. **In C**, representative three-dimensional reconstruction images of diffuse light imaging tomography are shown following three days of protocol. A second locus of intense TLR2 expression is present in mice exposed to IH only, and it co-localises to the region of brain’s septal nuclei, more so to medial septal nucleus. **D.** Photographs of a single mouse taken at different times during intermittent hypoxia show different sites of TLR2 induction in the brain (**D**). The colour calibrations at the right are photon counts. Note the occurrence of different scales in various ranges. *Abbreviations*: C- control; TLR2- Toll like receptor 2; IH- intermittent hypoxia; SN- medial septal nuclei; SD- standard deviation; OB –olfactory bulb.

Using this relatively novel *in vivo* imaging approach, we demonstrated the inflammatory response with a marked chronic component. As shown in Figure 1A, the photon emission and TLR2 signals/microglial activation were significantly elevated, when compared to control baseline levels, throughout a three-week period interval following a daily IH experimental protocol. The quantitative analysis of photon emissions revealed that levels of TLR2-induction signals peaked after 24 hours (1.16×10^7^±3.43×10^6^ p/s/cm^2^/sr, n=18) reaching statistical significance following three days of the protocol (*P*=.018; baseline: 7.68×106±1.81×106 p/s/cm^2^/sr, n= 20 *vs* day 3:1.12×107±2.40×106,n=17) (Figure 1A). A later smaller TLR2 peak was also noted following ten to twelve days of the experimental protocol. Thereafter, the total photon emission remained significantly elevated and relatively constant, up until the last analysed three-weeks timepoint (Figure1 A,D).

Olfactory bulb microglia in mice receive and translate numerous inputs from the brain and the environment and likely serve as sensors and/or modulators of brain inflammation. The subset of olfactory bulb microglial cells in mice was previously shown to continually express TLR2, enabling them to survey the environment in a ‘primed’ or alert state(19). Similarly, in our study, a baseline activation over the area of the olfactory bulb and anterior olfactory nucleus was recorded in both, control (TLR2CTRL) and IH (TLR2IH) mice (control 7.655×10^6^ps^−1^;n=6*vs*IH 7.821 × 10^6^ps^−1^;n=7). However, the TLR2 signal was stronger under IH and accompanied by an additional increase and widespread frontal distribution (Figures 1D, 2D-F). In addition, following 72 hours of IH protocol a significant activation in the region of the basal forebrain was recorded (Figures 1C; 2D-F), and then further traced to the micro-region of the septal nuclei via the 3D *ex vivo* reconstruction of the signal (Figure 2).

**Figure 2.**
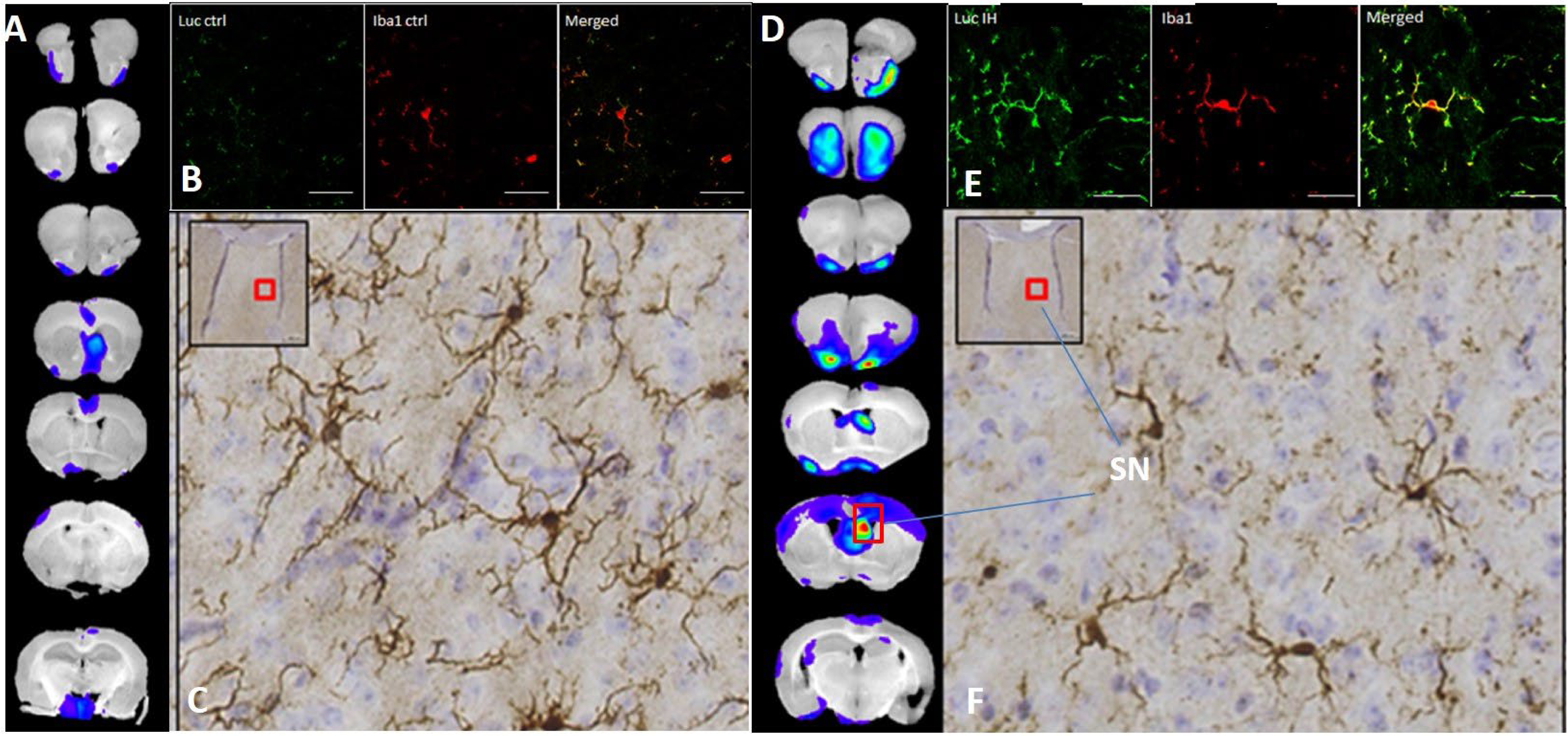
*Ex vivo* imaging of bioluminescence signal at 72 hours of experimental protocol co-localises induction of TLR2 microglial signal to basal forebrain and septal nuclei. Representative images of control (**A-C**) and animals exposed to IH (**D-F**) at 72 hours. All animals were exposed to *ex vivo* bioluminescence imaging where TLR2 signal was confirmed in olfactory bulb and anterior olfactory nucleus in both groups, whilst additional signal was noted in the septal nuclei (*red rectangle*) of the IH group only. Immunofluoresecent images of septal region in control (**B**) and IH animals (**E**) demonstrated colocalization (blue arrows) of luciferase (*luc*; green) and microglial marker (*Iba1*; red). In **F**, a distinct amoeboid activated morphology (blue arrow) of Iba1 positive cells in the septal nuclei region of IH animals is shown. Respective areas in red rectangles depict regions from which photomicrographs were taken. *(X40, Zeiss LSM 510 Meta confocal microscope; scale bar denotes 50 µm.)* *Abbreviations*: Ctrl- control; TLR2-Toll like receptor 2; IH-intermittent hypoxia, SN- septal nuclei.

### TLR2-transgene induction in the brain co-localises with microglia

To further characterize micro-regional differences of the inflammatory response recorded by BLI at the three key neuroanatomical sites - olfactory bulb and anterior olfactory nucleus (at baseline), septal nuclei (72 hours), and at later points, at the wider frontal cortical regions (see Figure 1) - we then explored cellular aetiology of TLR2-signal in TLR2CTRL and TLR2IH-mice (Figure 2). We were able to confirm that the endogenous TLR2-protein was indeed induced within these areas, and that the vast majority of cells expressing these receptors were microglia (Figure 2E).(19). As shown in larger magnification photomicrograph in Figure 2F, the luciferase immunoreactivity co-localized with *Iba1* immunostaining (microglial cells with amoeboid activated morphology) in the micro-region of septal nuclei, further suggesting the importance of this site early on in the inflammatory cascade.

### TLR2 modulates the effects of chronic IH on structural brain changes

In order to investigate whether initially demonstrated neuroinflammatory response later on resulted in structural plastic changes, known to occur in patients with OSA(2), we utilised high resolution *ex vivo* magnetic resonance imaging (MRI). To fully verify involvement of the TLR2 system, mice with and without (TLR2^−/−^) functional TLR2 gene were imaged after three weeks of IH or CTRL protocol (Figures1,2).

As shown in Figure 3, comparison of structural brain grey and white matter changes with MRI in TLR2IH vs CTRL mice demonstrated coexistent hyper- (*green*) and hypotrophic (*red*) cortical, subcortical and white matter changes. Most important enlargements were visible bilaterally in the hippocampi and presubiculi regions while the reduction of volume was most evident bilaterally in the reticular nuclei of the thalamus, dorsal striatum, parahippocampal and piriform cortex and dorsolateral pons, with involvement of distinct parts of periaqueductal grey (PAG), including the dorsal raphe nuclei (DRN) (Figure 3, also see FigureS2-4).

**Figure 3.**
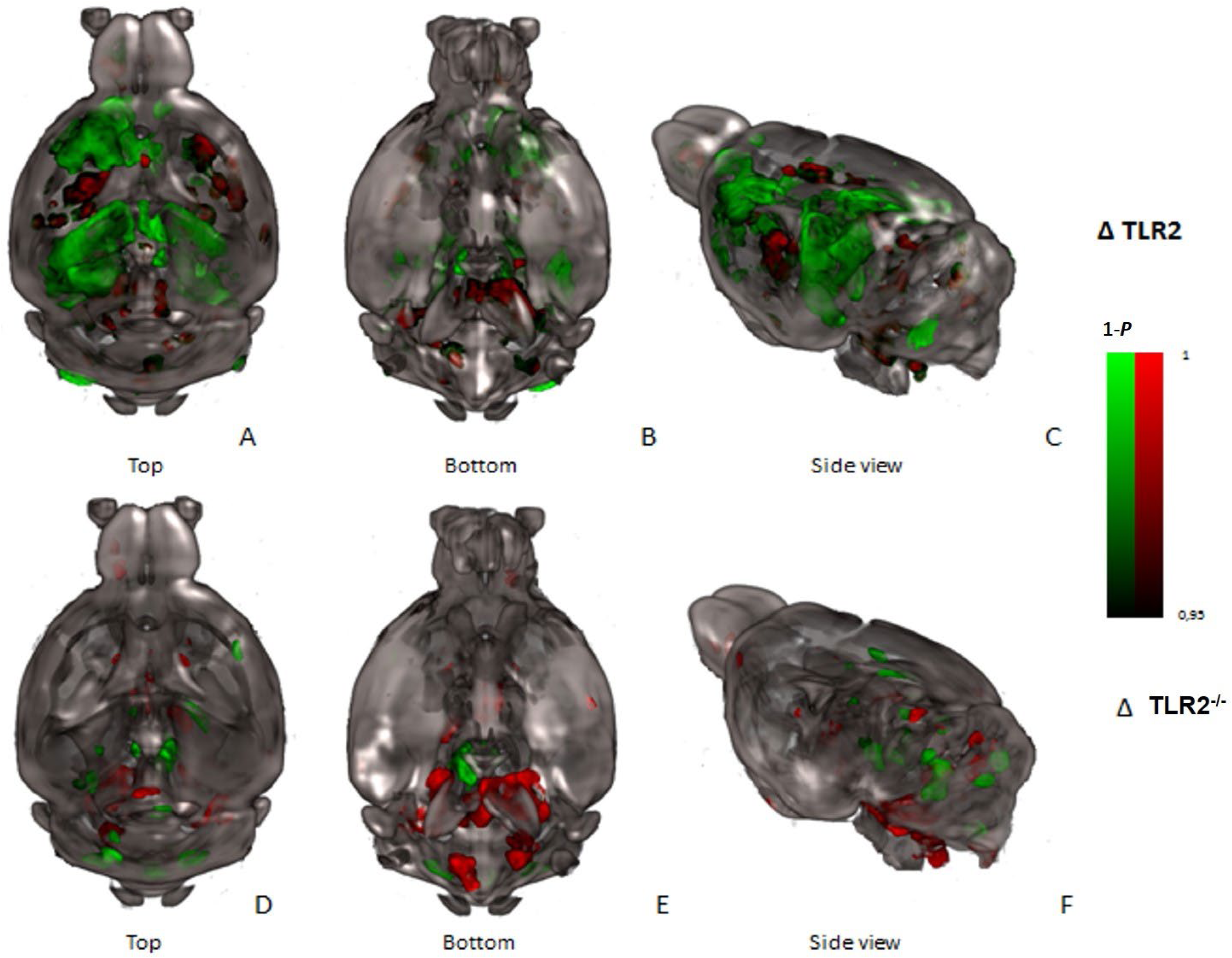
Three-dimensional rendering of all significant volume changes following three weeks of intermittent hypoxia protocol in mice with (TLR2: top row) and without TLR2 receptors (TLR2^−/−^:bottom row). Co-existent hyper- (*green* clusters) and hypotrophic differences (*red* clusters) in the mouse brains are demonstrated. Shown contours are statistically significant differences, i.e. intermittent hypoxia (IH) > control (C); *P*< .01. Data is displayed on the mouse template image(49) (50). *3D* reconstructions are shown from 3 perspectives: (**A, D**) Top view. (**B, E**) Bottom view. (**C, F**) Side view. *Hippocampus* is enlarged after IH as well as left part of the *motor* and *cingulate cortex* and *septum*, while these changes are not visible in TLR2^−/−^ mice. Suggested reductions of volume after IH are visible in the *reticular nuclei* of the *thalamus* and *dorsal striatum* of TLR2 IH and pronounced in *pons*, *medulla* and *mesencephalon* of TLR2^−/−^ IH. *Abbreviations*: 3D- three-dimensional; TLR2-Toll like receptor 2; IH-intermittent hypoxia.

Taken together, the spatio-temporal nature of the demonstrated neuroinflammatory process over the three weeks suggested that majority of later structural changes developed in neuroanatomical regions with monosynaptic connections to initial frontal and basal forebrain cortical sites of microglial TLR2 response (Figures 3,6). Conversely, significantly weaker and predominantly hypotrophic changes were shown in TLR2^−/−^ mice (Figure 3). Here, changes after three weeks were located in more posterior and inferior brain regions of mesencephalon, pons and medulla, including important sleep regulating brainstem structures of locus coeruleus, parafacial, parabrachial, PAG, dorsal raphe and pedunculopontine nuclei (Figures 3,S4).

**Figure 4.**
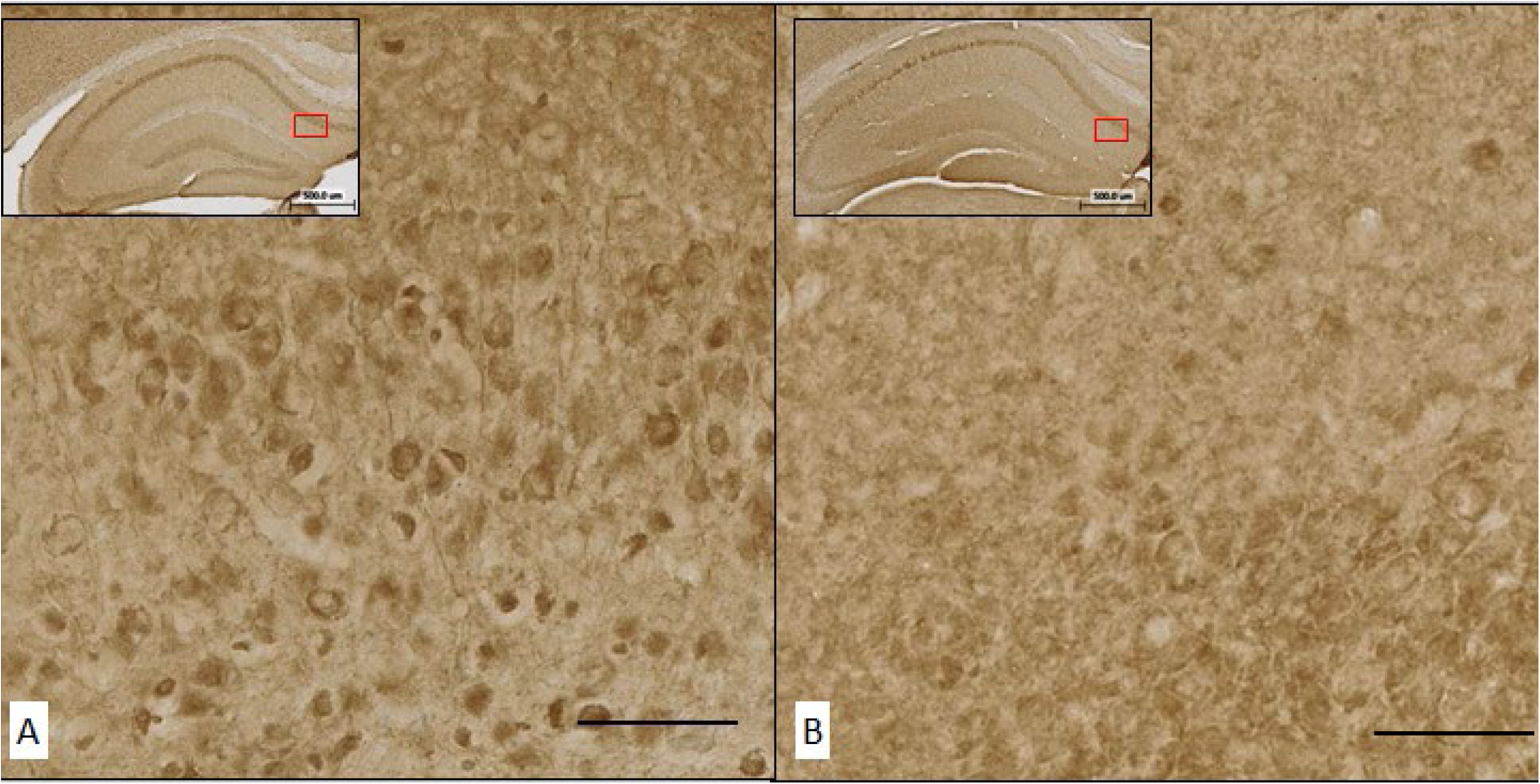
Representative images of *cFos* staining in the hippocampus following three weeks of intermittent hypoxia protocol in mice with (A) and without functional TLR2 (B). Digital stereological analysis revealed significantly higher number of *cFos* positive neurons in the group with functional TLR2 (**A**) than in the TLR2^−/−^ group (**B**) (*P*= .048). Shown are representative photomicrographs (40x; *BrainChopper*); red rectangles show the hippocampal region from which that the photomicrograph was taken from. Scale bar denotes 50 µm. *Abbreviations*: IH-intermittent hypoxia, TLR2-Toll like receptor 2.

**Figure 5.**
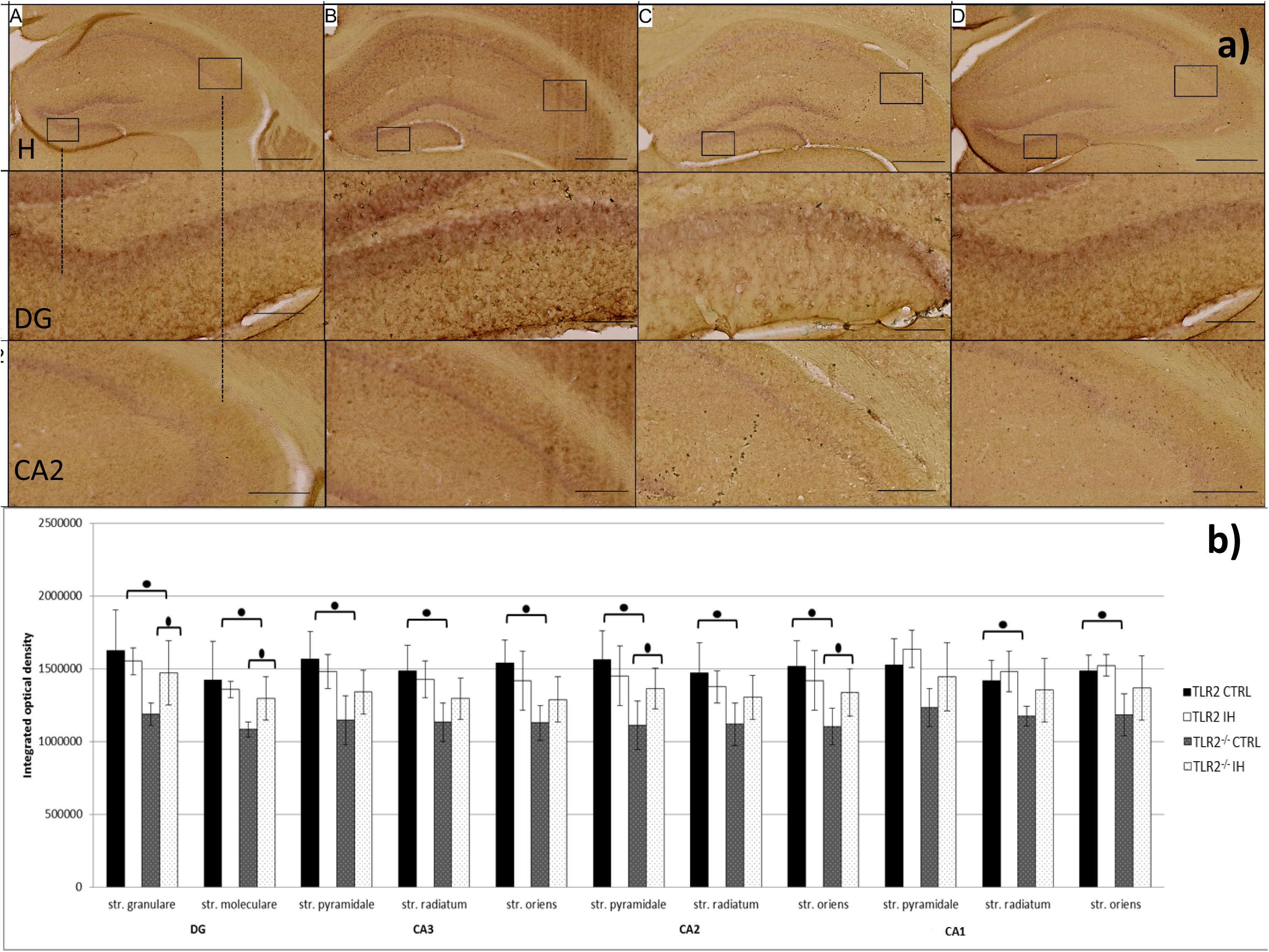
Representative images of neuroplastin staining in the hippocampus (a). Quantification of integrated optical density of showed significantly lower neuroplastin immunoreactivity in TLR2^−/−^ CTRL (**C**) animals compared to TLR2 (**A**) in almost all hippocampal regions. Scale bar in H (5x) denotes 500 μm; in DG (20x) denotes 100 μm; in CA2 (10x) denotes 250 μm. **Quantification of integrated optical density shows significantly higher neuroplastin immunoreactivity in animals with functional TLR2 systems in almost all major hippocampal regions (b).** Also, quantification showed increase of neuroplastin immunoreactivity in TLR2^−/−^ animals exposed to IH protocol in *dentate gyrus* (DG) *stratum granulare* (*P*=.023) and *stratum moleculare* (*P*=.013), and in CA2 region *stratum pyramidale* (*P*=.024) and *stratum oriens* (*P*=.028). Error bars denote standard deviation (SD). * equals *P* < .05, Student t-test. *Abbreviations*: CA- Cornu Ammonis; CTRL – control conditions; DG – dentate gyrus; H-hippocampus; IH-intermittent hypoxia protocol, TLR2-Toll like receptor 2.

**Figure 6.**
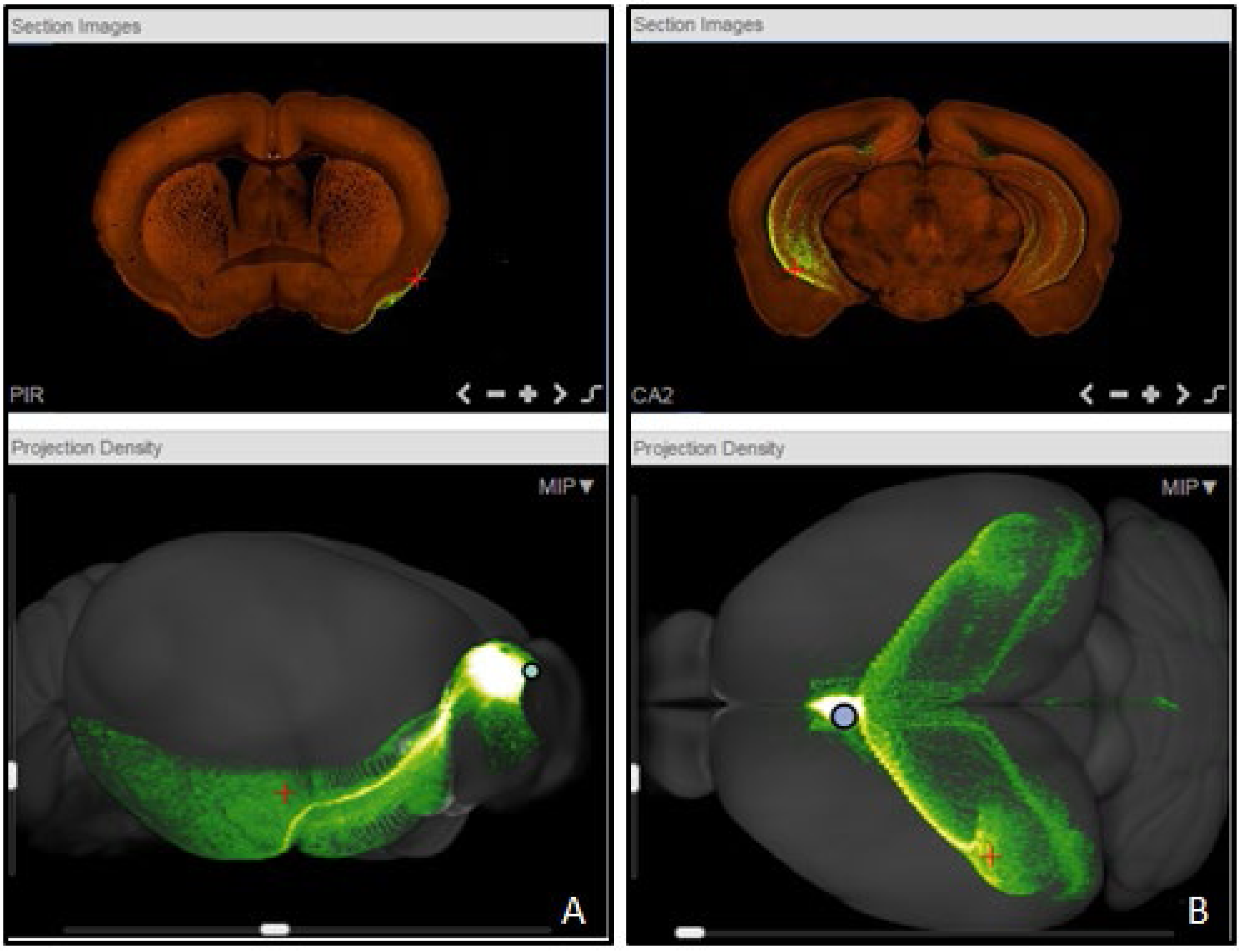
Representative coronal sections (first row) depict injection sites of rAAV tracer into the olfactory bulb (A), and septal nuclei (B) respectively (adapted from Allen Brain Atlas(51)). 3D tracing (second row) of distinct monosynaptic pathways from injected two regions is shown. **A:** monosynaptic pathways from OB are visualised; pathways can be visualised to traverse posteriorly and extend to the entorhinal cortex (*right view*). **B**: septal nuclei’ monosynaptic projections innervate wide bilateral hippocampi regions (*top view*).(*51*) Of note is that the affected anatomical regions highly correspond to neuroanatomical structural changes observed following three weeks of our sleep apnoea model in mice with functional TLR2. *Abbreviations*: retrograde labelling with adeno-associated virus (rAAV), TLR2-Toll like receptor 2.

Further immunochemistry analyses examined the ultrastructural aetiology of observed neuroimaging findings. The TLR2 genotype appeared protective against demyelinating effects of IH. For example, a widespread IH-induced demyelination of the hippocampus was only demonstrable in TLR2 deficient mice (Figure S7;*P*_TLR2IH*vs* TLR2CTRL_=.0286), whilst it did not reach statistical significance in mice with functional TLR2 system (Figure S6). However, despite distinct morphological and cytoskeletal differences between astrocytes and microglia in four investigated groups, no statistically quantifiable changes in numbers of astroglia cells were recorded in respective regions of interest (ROIs) (Figures S5,6; Supplement). Potential molecular drivers of noted enlargements in the ROIs in TLR2IH and TLR^−/−^IH mice were then further explored.

A prominent up-regulation of immediate early gene c-*fos* was recorded in the hippocampal regions of the cerebral cortex in mice with functional TLR2-system (Figure 4). Similarly, we demonstrated a more prominent up-regulation of the cell adhesion molecule neuroplastin in mice with a functional TLR2-system, as shown by its stronger immunoreactivity signal pattern in all major hippocampal sublayers that contain neurons (e.g. granular layer in dentate gyrus (DG), and pyramidal layers in *Cornu Ammonis* (CA)) (Figure 5). However, of note is that under our experimental conditions, the TLR2 deficiency appeared permissive for a more prominent modulation of IH-induced inflammatory stress response via neuroplastin in distinct hippocampal DG *stratum granulare* (*P*=.023) and *moleculare* (*P*=.013), and in CA2 region *stratum pyramidale* (*P*=.024) and *stratum oriens* (*P*=.028)(Figure 5, Supplemental Table S3).

### TLR2-induced neuroinflammatory plastic responses localised to brain derived neurotrophic factor (BDNF), neuroplastin and fibronectin-1 (FN1)-rich neurocircuitry

Together, our data provides further evidence for the role of TLR2 in modulation of the neuroinflammatory response, and suggested its spatio-temporal spread via synaptically connected enthorhinal and basal forebrain networks.

We further examined the molecular origins underlying neuromechanisms behind OSA-induced cortical network reorganization. To this end, an investigation of the observed network’s modulation in mice with and without functional TLR2 was undertaken by linking it with the microregional brain plasticity gene expression profiles (Figure 3). For this purpose the *Allen Brain Mouse Atlas*(*22*) mRNA gene profile expression database and MR parametric *t*-statistic maps were utilised via a novel mapping method, recently described(23) (Figures 7, also see Table S5). A modulatory role in observed structural changes was suggested for several major neuroplasticity genes, however, this association was evident only in mice with a functional TLR2-system (Table S4). The genes that were linked with TLR2-enabled structural network reorganisations were the brain derived neurotrophic factor (BDNF), neuroplastin, Calcium/calmodulin dependent protein kinase II alpha (CAMK2A), Rho Guanine Nucleotide Exchange Factor 6 (ARHGEF6), Cholecystokinin (CCK), Fibronectin 1 (FN1) and RAS guanyl nucleotide-releasing protein 1 (RASGRP1)(TableS4). Further multiple regression analysis suggested the strongest, statistically significant permissive role for BDNF, FN1 and RASGRP1. A significant correlation of longitudinal relaxation time (T1) with BDNF expression was shown (r=0.819;*P*=.002) (Figure 8). FN1gene expression was predictive of brain volume increases (r=0.627;*P*=.039). Conversely, RASGRP1 gene had an inverse relationship to transverse relaxation time (T2, r=-.646;*P*=.033) (Figure8), suggestive of its protective microregional role in topical inflammation. Strikingly, we again failed to demonstrate link between any of the investigated neuroplasticity genes in TLR2 deficient mice (TLR2^−/−^) with any of their inflammation-induced structural changes.

**Figure 7.**
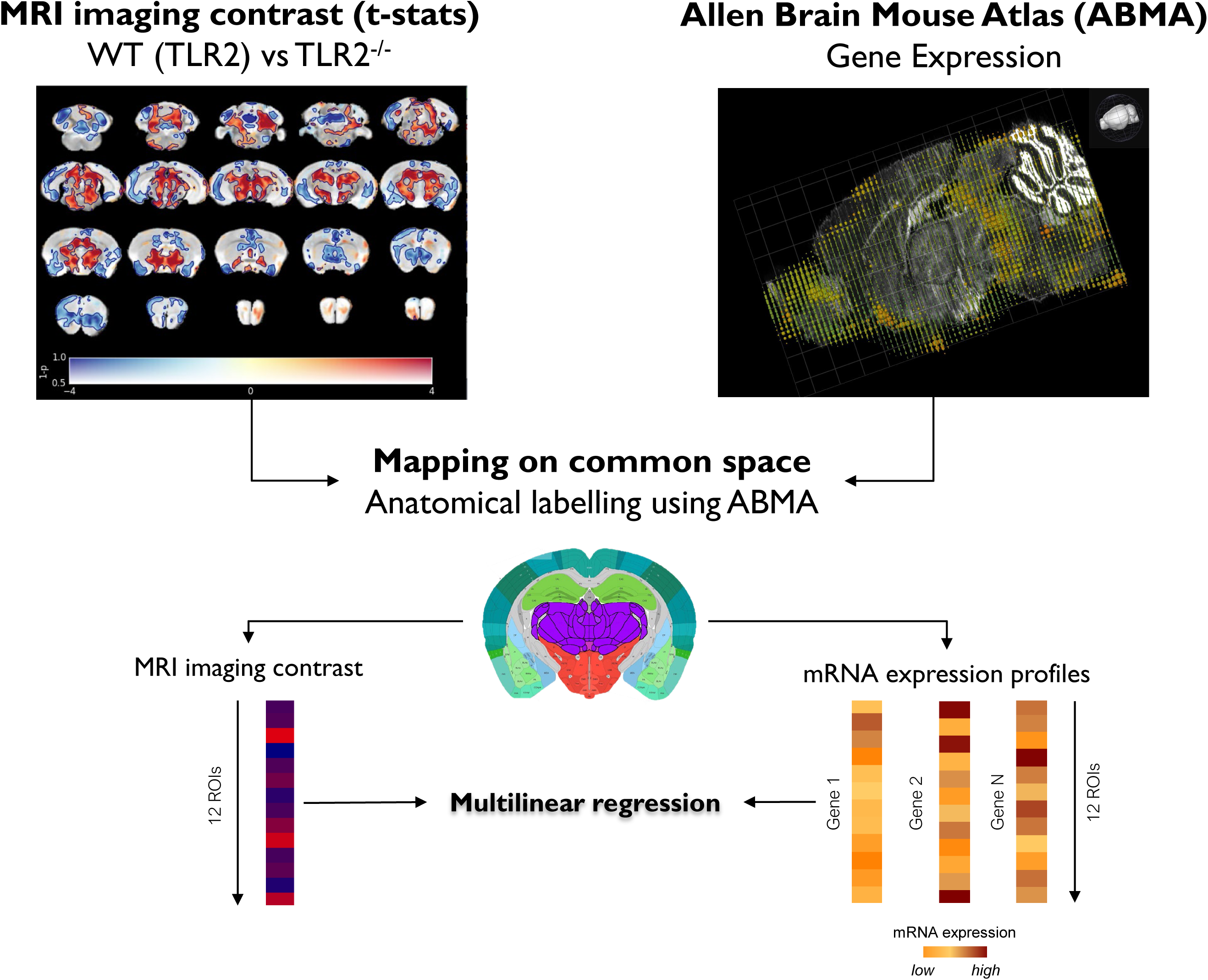
Schematic presentation of the MR-gene mapping research protocol: imaging *versus* mRNA expression analysis pipeline is shown. Effect of genotype (shown *top left*): voxel-wise differences in brain volume between TRL2 and TLR2^−/−^ mice. Data are shown as dual-coded statistical maps in which volume change is coded by colour hue, and the family-wise error (FWE) corrected *P*-values are coded by transparency. Warm (red) colours indicate larger volume in TLR2 *^−/−^* mice. Areas in which FWE-corrected *P* < .05 are contoured in black. Non-parametric statistics were performed using FSL randomize with 5000 permutations and threshold-free cluster enhancement.

**Figure 8.**
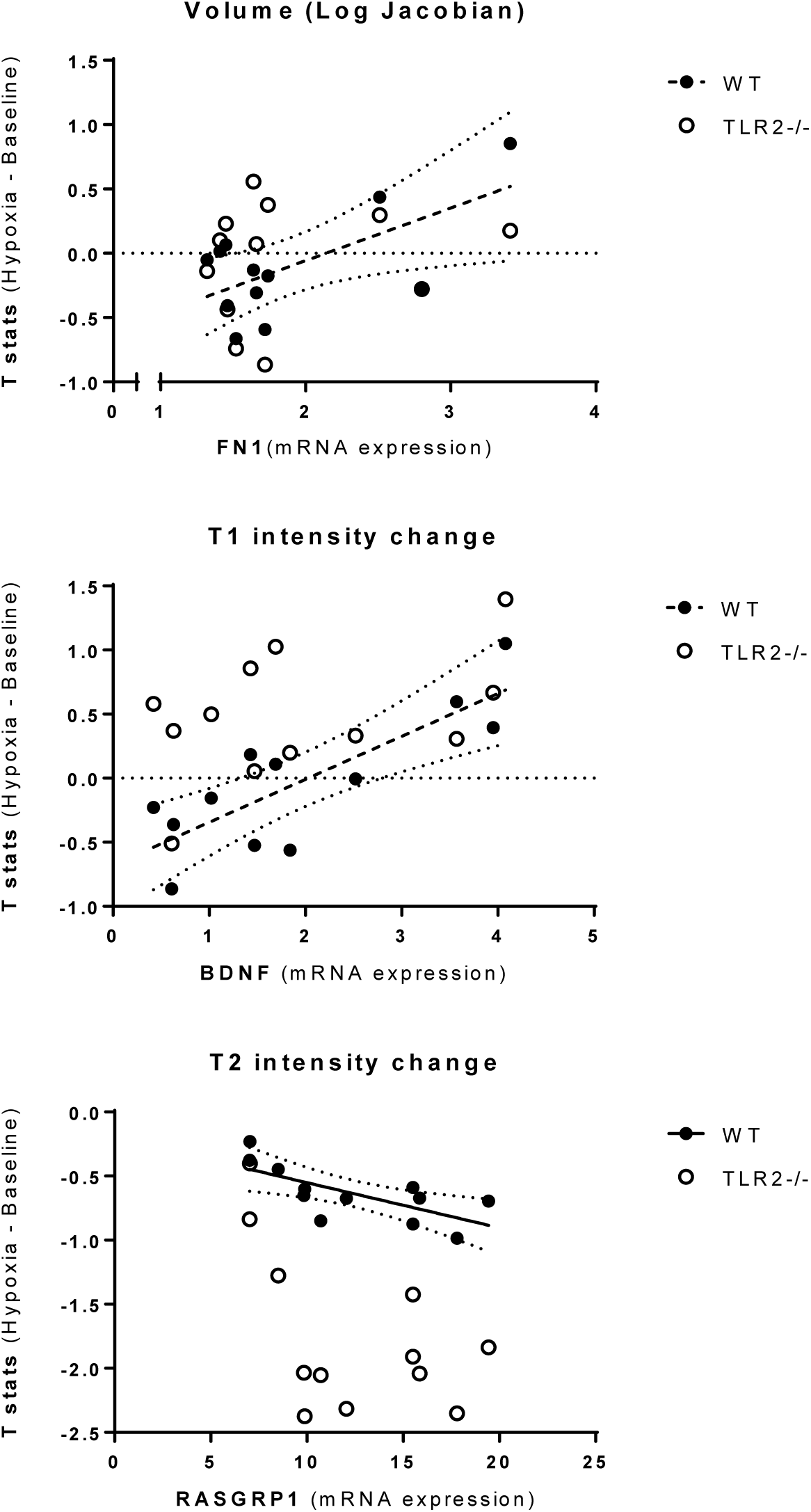
MRI imaging and gene expression correlation. All the correlations with wild type mice (WT) were found to be statistically significant. All the correlation with TLR2^−/−^ were reported as not significant. *Abbreviations*: ABMA, Allen Brain Mice Atlas; TLR2-Toll like receptor 2.

### Neuroinflammatory response: from effects on mood, cognition to effects on weight gain?

Finally, we wanted to assess if mice in our OSA model shared similar functional and behavioural changes to those frequently reported in patients with OSA, including their well-documented struggles with obesity, somnolence, fragile mood and memory and attention deficits(2, 24). We also wanted to see if those changes were linked to neuroanatomical ROIs that were initially highlighted by our BL and MR imaging findings: frontal cortex(16), septal nuclei, ventral hippocampi(25) and PAG(26, 27).

*Weight*: Firstly, the adaptive role for TLR2 system in weight gain was investigated. A newly proposed anorexigenic neural circuitry in rodents incorporates two ROIs implicated in the neuroinflammatory response in our study, the ventral hippocampus and lateral septal nucleus in the brain.(25) A set of structural comparisons of ROIs was done and significant differences demonstrated in the volumes of lateral septal nuclei (V(%):TLR2IH 0.373±0.007*vs* TLR2^−/−^IH 0.335±0.004;*P*=.01) and in ventral hippocampi (V(%):TLR2IH 0.0028±0.0001*vs* TLR2^−/−^IH 0.00294±0.00004;*P*=.0002) possibly suggesting functional TLR2-driven adaptive value of noted changes in mice exposed to IH. Similar structural differences were noted in control groups, although they were statistically remarkable only in the anorexigenic region of the right ventral hippocampi (V(%):TLR2CTRL 0.00273±0.00004*vs*TLR2^−/−^CTRL 0.00308±0.00007;*P*=.0001). To further test if there is a stronger baseline anorexigenic structural component in the neural circuit of TLR2 deficient knockout mice, we investigated the effect of TLR2 genotype on the weight gain. In keeping with our structural data, a strong link between a functional TLR2 system and an adaptive ability to gain weight was demonstrated, both under control (TLR2CTRL:third day(%): 1.17±2.86;sixth day:3.58±2.39;ninth day:6.25±10.14;16^th^ day:7.64±4.54;20^th^ day: 5.60± 4.34) and IH experimental conditions (TLR2IH:third day(%):-6.28 ± 5.43; sixth day:-4.43 ± 4.49; ninth day:-4.31±5.04;16^th^day:-3.35±6.1;20^th^day:-1.99±6.05)(Figure 9). TLR2^−/−^ deficient mice (TLR2^−/−^CTRL), on the other hand, failed to gain further weight over the period of three weeks, and their weight remained constant throughout the protocol (Figure 9). Over a three-week period, a significant weight loss in both TLR2IH and TLR2^−/−^IH mice was recorded, compared to their respective controls (Figure 9, Supplement). This reached a statistical significance after three days, and it remained significant until the end of the study (Figure 9;Tables S7,8). However, whilst mice with functional TLR2 showed a steady state increase in weight following the initial loss, TLR2^−/−^ mice were not able to regain initially lost weight (Figure 9).

**Figure 9.**
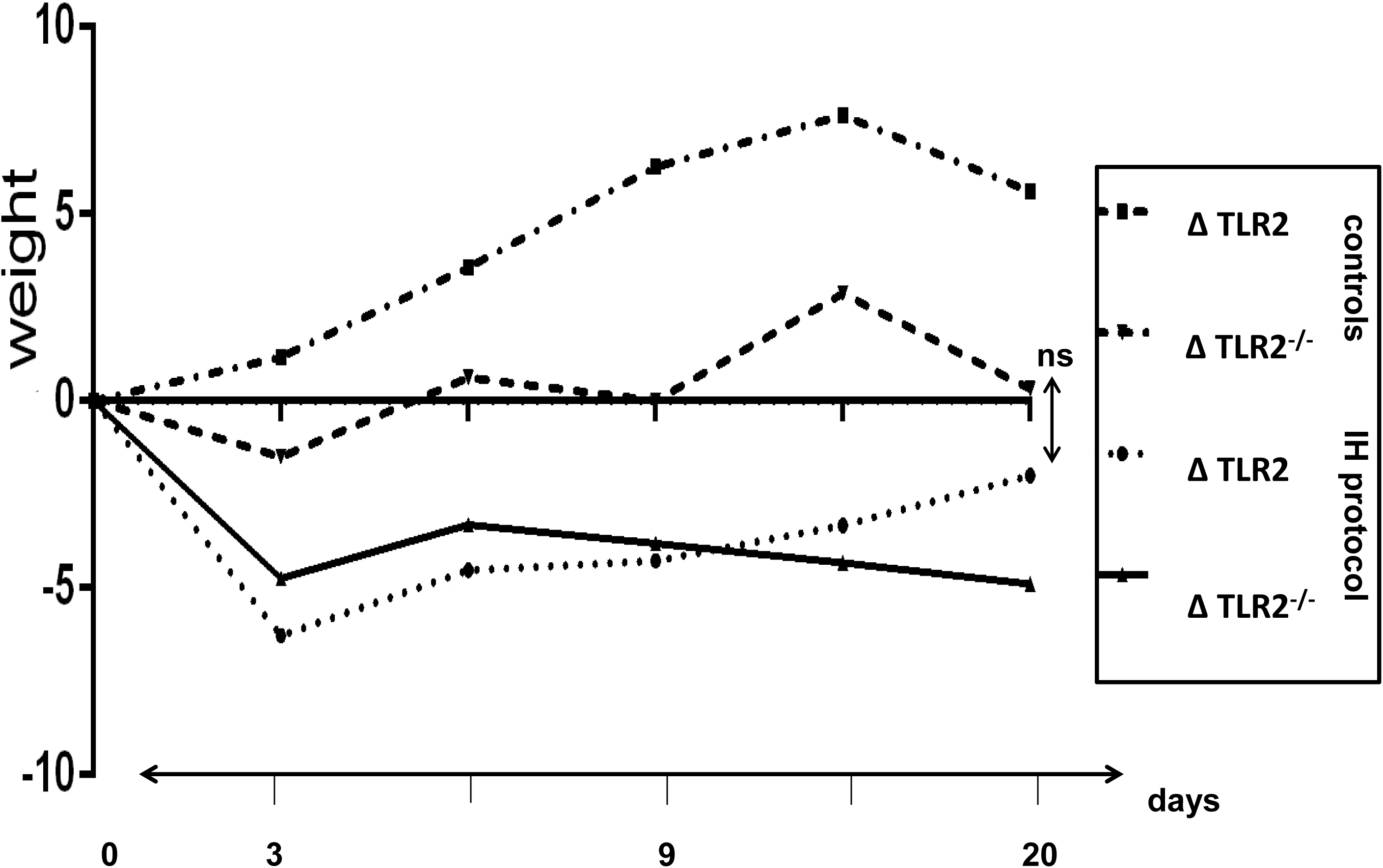
Significant body weight loss induced by intermittent hypoxia. Weight changes over the course of three weeks of experimental conditions in 4 investigated groups are shown: two groups with functional TLR2 (CTRL-control or IH-intermittent hypoxia protocol) and two groups with knockout TLR2^−/−^ (CTRL-control or IH-intermittent hypoxia protocol). IH protocol was associated with significant weight loss after three days of protocol. However, a compensation of weight loss through time occurred in mice with functional TLR2 gene, while weight loss recovery was not recorded in knockout mice (TLR2^−/−^). Overall, mice with a functional TLR2 gene in control group showed steady rise in weight while knockout mice keep a steady weight throughout that time period. *Y axis* denotes time points of 0, 3, 5, 9, 15 and 20 days of experimental protocol. Values ±SD and statistical significance are presented in tables S7 and S8 of the Supplement. *Abbreviations*: CTRL-control; IH – intermittent hypoxia protocol; SD- standard deviation, TLR2-Toll like receptor 2.

*Mood and cognition*: Next, the adaptive role for TLR2 system in neurocognitive changes was investigated. Using several behavioural tests (Table S6) known to assess and target psychomotor changes, affective and cognitive symptoms, we observed two primary findings, which were consistently replicated across several behavioural tests and their parameters (Table S6). Firstly, as expected, we demonstrated increased psychomotoric agitation and anxiety in all mice exposed to the IH protocol. This behaviour appeared independent of their TLR2-functionality. For example, during the tail suspension test (TST) the mice that were exposed to the experimental IH protocol spent significantly more time trying to escape the uncomfortable setting, and less time being immobile (immobility period:TLR2IH:103.72±48.09s;n=15;TLR2^−/−^IH:99.43±33.17s;n=16) compared to their respective control groups (TLR2CTRL: *P*=.051; 152.68±34.69s,n=12; TLR2^−/−^CTRL: *P*=.014; 144.90±56.08s,n=8).

The second finding was unexpected. It was noted that functional TLR2 genotype in mice exposed to IH protocol was linked to a specific proactive adaptive behavioural endophenotype. This was, for example, demonstrated by shorter latencies to trying to first escape during the TST (TST: TLR2IH27.65±7.11s;TLR2CTRL 27.37±12.79s *versus* TLR2^−/−^IH 45.04±22.08s;TLR2^−/−^CTRL 30.14±12.38s). Here functional TLR2 genotype appeared to rescue depression-like behaviour noted under experimental IH conditions of repeated stress (*P*_(TLR2IH*vs*TLR2^−/−^IH)_ =.027). We then investigated which ROIs might have functionally contributed to this behavioural endophenotype in TLR2IH mice. Correlation analyses suggested significant divergent link to (left) frontal cortical (*P*=.02,r=.635) and periaqueductal grey region (*P*=.083,r=-.499; also see Table S1). A significant aberrant connectivity between the two regions was also noted (*P*=.011;r=-.66).

In regard to other behavioural findings, only a statistical trend for more effective spatial navigation and cognitive processing was seen in mice with functional TLR2. For example, in Y-maze test the presence of TLR2 system appeared to partially rescue deficit in spatial acquisition following exposure to IH, otherwise recorded in TLR2^−/−^ mice, as measured by the path efficiency (path efficiency%: TLR2IH 1.05±0.16; TLR2CTRL 1.09±0.20;TLR2^−/−^IH 0.63±0.14;TLR2^−/−^CTRL 1.04±0.20; *P*_(TLR2IHvsTLR2^−/−^IH)_=.06). (see Table S6 for further details). Several other differential modulatory trends of genotype and phenotype on behavioural and cognitive parameters of mice are presented in Figure 10.

**Figure 10.**
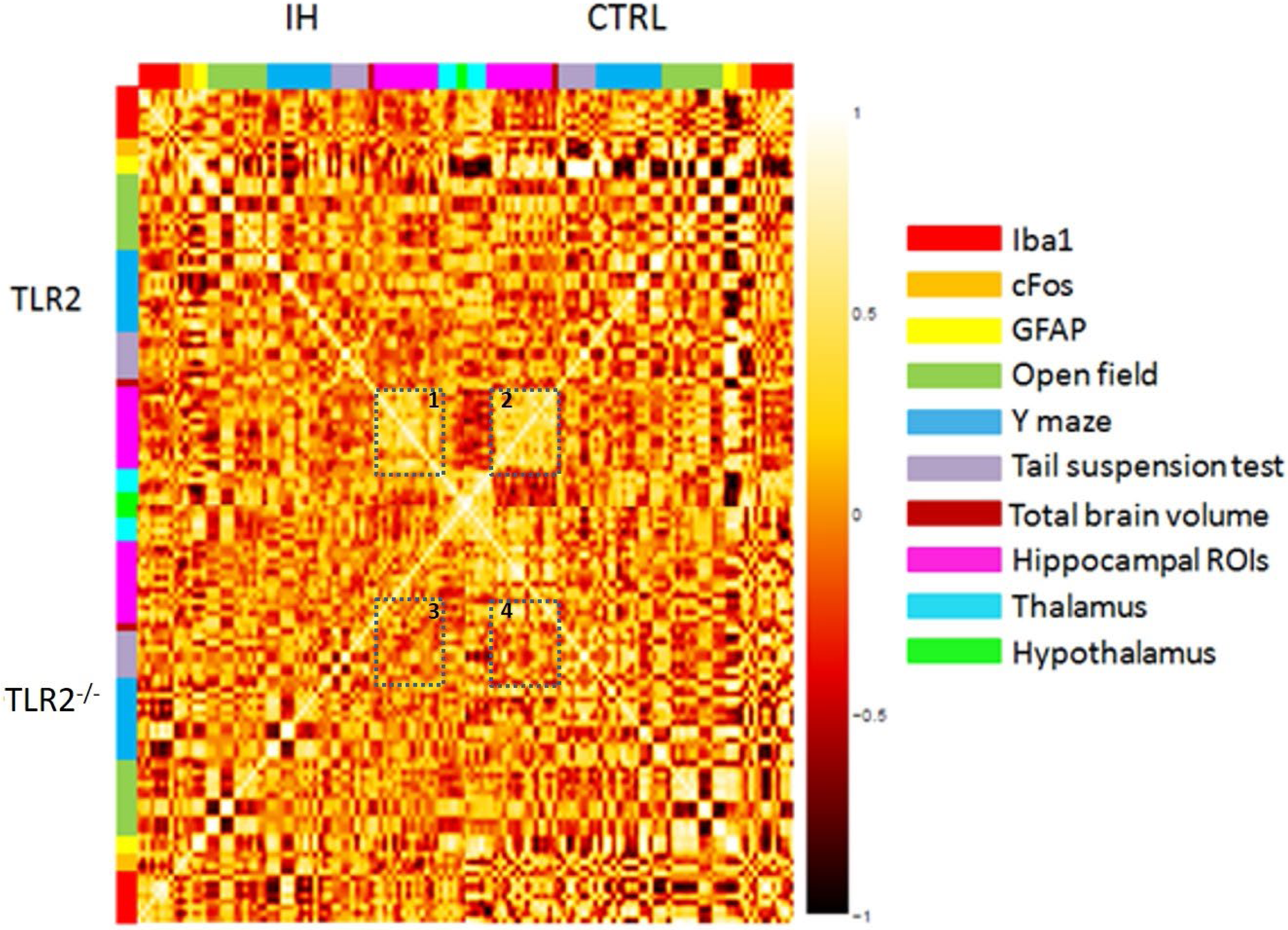
Heat map of selected *post hoc* correlational analyses of histological, behaviour and bilateral structural MRI findings. Plastic changes are visualised through a heat map presentation of the Pearson’s correlation values, with the *top row* showing results for mice with functional TLR2 system (e.g. **1**-TLR2 IH and **2**-TLR2 CTRL), and the *bottom row* showing mirror image of results for the knock out mice (e.g. **3**-TLR2^−/−^IH and **4**-TLR2^−/−^ CTRL). The superimposed “square shapes” are placed over values for the hippocampal ROIs, and demonstrate strong positive bilateral structural links in TLR2 control mice (see **2**; TLR2 CTRL), with “square” appearing less focused and more ambiguous in mice exposed to IH conditions (see **1**; TLR2IH). The clear shape boundaries appear further diffused in mice without functional TLR2 (see **3 and 4;** TLR2^−/−^IH and TLR2^−/−^CTRL). Adaptive plastic changes in the hippocampal structures under control conditions in the mice without functional TLR2 system (see **4**; TLR−/− CTRL), might be suggestive of distinct neurodevelopmental adaptation in this region. Further loss of positive structural correlations shown in the knockout mice exposed to the IH protocol (see **3**; TLR2^−/−^ IH mice) in this region, on the other hand, might be suggestive of a differential inflammatory response to that instigated in 2, in mice with a functional TLR2 system. From upper and left side the colours denote grouped variables: *Iba1* (*red*), *cFos* (*orange*), *GFAP* (*yellow*), open field test (*green*), Y maze test (*blue*), tail suspension test (*gray*), overall brain volume (*maroon*), hippocampus (*lilac*), thalamus (*light blue*), hypothalamus (*light green*). The upper half of the map pertains to TLR2 groups (1 and 2) while the lower denotes TLR2^−/−^ groups (3 and 4). Results for groups exposed to IH protocol are in the left part of the heat map, while the control groups are on the right. The colour scale on the right matches colour hue with the value of the corrected Pearson’s correlation coefficient where white represents r=1 and dark red r=-1. *Abbreviations*: cFos- nuclear phosphoprotein; CTRL- control; GFAP- Glial fibrillary acidic protein; *iba1*-ionized calcium binding adaptor molecule 1; IH – intermittent hypoxia protocol; ROI- regions of interest; TLR2-Toll like receptor 2.

## Discussion

Our sets of results converge on a single conclusion, that OSA instigates an acute neuroinflammatory response in the basal forebrain, and more chronically in other parts of frontal cortex, which consequently leads to a true network-specific structural changes with functional and further metabolic alterations. TLR2 system-driven (mal)adaptive changes in fronto-brainstem and hippocampal-septal circuitry are demonstrated, with links to agitated (mal)adaptive behaviour under episodes of stress, and an increased ability to gain weight.

Patients with OSA are known to be prone to obesity(28), they have well documented cognitive and neuropsychiatric deficits(24), excessive daytime somnolence, and are more likely to develop depression and anxiety.(3, 24, 29) They are also known to be particularly prone to traffic and general work accidents, in part through their well-documented deficits in attention(24), but also presumably through the erroneous encoding of spatial information in the context of navigation.(2, 30) Somewhat surprisingly, our data suggested an early antidepressogenic effect of the neuroinflammatory response in OSA that is TLR2-dependent, and which is functionally linked to a distinct fronto-brainstem subcircuitry. The activation of a similar network in mice has been reported to favour effortful behavioural responses to challenging situations.(31) More specifically, it was shown that a selective activation of a subclass of prefrontal cortex cells, which project to the brainstem, has been able to induce a profound, rapid and reversible effect on selection of the active behavioural states(31). According to a description of traditional learned-helplessness response(32), depressed patients do not favour effortful behavioural responses to challenging situations. To the contrary they are, through their own overvalued negative evaluations of challenges, significantly more likely to earlier retire from trying to find solutions out of their predicament. Conversely, in our study, mice demonstrated a clear, at least initially adaptive, ‘hyperactive’ behavioural response when placed in the challenging situation. They, unlike their TLR2-deficient counterparts, continued to try to find solutions out of their predicament. It would be of paramount interest to see if this initial adaptive behavioural response with time develops into a potentially more dangerous, futile, and energy wasting ‘agitated’ depressive profile(32, 33), with heightened anxiety component. A similar mixed anxiety and depression endophenotype has indeed been previously described in some patients with OSA(34), and it has been traditionally linked with higher suicide risks in depressed patients(35). Strikingly, only recently, in postmortem brains of depressed patients who committed suicide, TLR2-protein and its mRNA gene expression have been shown significantly increased, especially in their frontal cortical regions(14). Hence, an early induction of the neuroinflammatory process via TLR2-system in frontal regions and basal forebrain, such as was demonstrated in our mice, may be of particular importance in understanding the neural circuitry underlying pathological behavioural patterns of action selection and motivation in behaviour of patients with OSA(3).

Our data suggests an adaptive side to the neuroinflammatory response in regards to feeding(33) and or ability to gain weight. OSA has been associated with significant metabolic changes, diabetes and weight gain, thought to be at least partly modulated through its systemic inflammatory effects(4). It has been previously proposed that a significant interplay between central nervous inflammation and systemic inflammation might occur in affected patients, which then further modulates bidirectional links between metabolism, weight gain and neurological disorders(12). More specifically, our results could be taken to propose that activated TLR2-system enables continued feeding drive under conditions of repeated stress (Figure 9). Presumably, similar drive could later cause maladaptive inability to control one’s food intake and to lead to an increased link between our emotional states and eating, causing an excessive weight gain in patients with OSA. The feeding circuitry that controls emotional or cognitive aspects of food intake is still largely unknown. However, recently, Sweeney and Yang(2015) demonstrated an anorexigenic neural circuit originating from ventral hippocampus to lateral septal nuclei in the brain, revealing a potential therapeutic target for the treatment of anorexia or other appetite disorders(25). Interestingly, we were able to differentiate this network solely based on the functionality of the TLR2-system in our mice, and severity of changes appeared further exacerbated by the IH stress (also see Table S1 for further details). Whilst preliminary, these findings might go some way to suggest an aberrant feeding circuit that controls emotional and cognitive aspects of food intake in patients with OSA.

In conclusion, utilising the rodent model of OSA that verifiably replicates arousals and hypoxaemia in patients with OSA(17) and visualising *in vivo* TLR2-activation, we were able to demonstrate early activation of microglia in regions of basal forebrain, with later widespread frontal projections, suggesting a pivotal role for TLR2 in brain’s response to OSA injury. Our results provide the first *in vivo* evidence of an OSA-induced inflammatory response, with septal nuclei, the major source of cholinergic input to the hippocampus, being highlighted as a region of particular vulnerability early on in the neuroinflammatory process (Figure 1C). Moreover, we were also able to link the initial neuroinflammatory response to later hypotrophic and hypertrophic neuroanatomical changes, with potential primary molecular drivers also highlighted by our results. Through these findings, a fingerprint of a distinct OSA-effected neurocircuitry has emerged, with frontal regions and septal nuclei being suggested as an initial TLR2-dependant seed sites.

Our neuroimaging results are in broad agreement with previous findings in patients with OSA where compensatory mechanisms, activation of various homeostatic gene programmes and astroglial neurogenesis have all been proposed to underlie some of the initial adaptive and later maladaptive changes(36, 37). In further agreement, our data also suggests that differential plastic response to OSA in the brain may depend on regional genes expressional profiles(38) For example our ‘MR-gene’ mapping findings support permissive and cohesive role for the TLR2-system in the interplay with BDNF, RSGRP1, fibronectin and neuroplastin-driven significant discrete and transformative neurophysiologic and behavioural changes.

Of the listed genes, the enhanced BDNF-levels have long been associated with increased plasticity and promotion of a growth-permissive environment, and its depletion with many neurological and psychiatric disorders(39). The concept of the interplay between BDNF and TLR2-system in our study might also explain proadaptive (preconditioning) protective effect on visuo-spatial memory in our TLR2IH mice, which, however, showed only significant trend. BDNF has been shown to enhance synaptic plasticity and neuron function in response to physical activity, learning and memory, and its baseline expression and activity-dependent upregulation in the hippocampus is believed to be under control of the medial septal nuclei, and important ROI in our study, with an important regulatory role in REM sleep (Figure 2). Patients with OSA show a rapid decrease in serum and plasma BDNF levels during initiation of the treatment (with PAP-device), likely reflecting enhanced neuronal demand for BDNF in this condition.(40) Similarly, OSA patients had increased TLR2-expressions on blood immune cells, which could be reversed with PAP treatment(41). TLR2-deficiency has been shown to impair neurogenesis previously(42) and more recently, the TLR2-receptor has been shown to enhance adult neurogenesis in the hippocampal DG after cerebral ischaemia(43). Of particular note to our findings, the loss of TLR2 has been recently shown to abolish repeated social defeat stress-induced social avoidance and anxiety in mice, and its deficiency mitigated stress-induced neuronal response attenuation, dendritic atrophy, and microglial activation in the medial prefrontal cortex(16). TLR2 has also been shown to modulate inflammatory response caused by cerebral ischemia and reperfusion via linking to endogenous ligands, such as fibronectin(44). Fibronectin, on the other hand, has been shown to act as a reparative molecule that promotes cellular growth and its levels are enhanced after brain injury in variety of disorders, including the AD(39, 45). Finally, neuroplastin is known to play a role in synaptic plasticity (e.g. long-term potentiation), formation and a balance of the excitatory/inhibitory synapses(46), in shaping of brain’s cortical thickness(47), and its involvement in early tissue response in hippocampi of AD patients has also been recently shown(48).

Taken together, the implicated gene programmes also indirectly suggest that modulations in neurites, cytoskeletal and receptor signalling, cell adhesion, axonal sprouting and other extracellular and perineuronal nets likely underlie the observed structural and functional changes. Whilst our findings are striking and theory-forming, our study leaves many questions still unanswered. There are several limitations to our findings behind OSA-injury, notwithstanding that the majority of experiments were done in cross-sectional manner, preventing us from deduction of causality and or direction of noted changes. Ideally, multimodal longitudinal *in vivo* manipulation of the microglia TLR2 in the highlighted ROIs, along with *in vivo* functional MR and further behavioural, genetic and electroencephalographic studies, should help shed much needed insight into here proposed novel neural mechanisms. Nonetheless, we believe that the distinct regional association between highlighted gene profiles and structural and functional changes, as demonstrated by our data, arguably further indicates a true circuitry-specific, rather than wider systemic, nature of TLR2-modulated neuroinflammatory injury and as such provide a plethora of potential investigational targets for future studies.

## Acknowledgments

This work was supported by the Wellcome Trust [103952/Z/14/Z]. M.V. was supported by National Institute for Health Research (NIHR) Biomedical Research Centre at South London and Maudsley NHS Foundation Trust and KCL. This work was further supported by the following projects: YoungBrain (ESF, HR.3.2.01) and Orastem (IP-2016-06-9451), awarded by Croatian National Foundation to D.M., and the following grants: EU FP7 grant GlowBrain (REGPOT–2012–CT2012–316120), EU European Regional Development Fund, Operational Programme Competitiveness and Cohesion, grant agreement No.KK.01.1.1.01.0007, CoRE – Neuro, and by Croatian Science Foundation project IP-06-2016-1892. We are grateful for all discussions, kind help and materials from the following colleagues and collaborators: Jasna Kriz, Raphaella Winsky-Sommerer, Paul Francis and William Crum. We are additionally particularly grateful to Jasna Kriz for providing Tlr2-luc/gfp transgenic mice, and to GlowLab multimodal imaging facility, University of Zagreb School of Medicine, Zagreb, Croatia, where the BLI scans were performed. This work is dedicated in memoriam of an esteemed Croatian scientist, neuropsychiatrist and a patriot, a dear friend gone too soon, Professor Vlado Jukic (Osoje, 1951-Zagreb, 2019).

## Contributors

All authors contributed equally to the writing and revision of this manuscript.

The authors do not have any conflicts of interest to report. The authors apologize to all the colleagues whose outstanding work could not be cited due to space limitations.

