## Supplement for "The Innate Immune Toll-Like Receptor-2 modulates the Depressogenic and Anorexiolytic Neuroinflammatory Response in Obstructive Sleep Apnoea"

**Running Title**: Sleep Apnoea and Neuroinflammation

### Supplement Outline

1. Materials and Methods

- Figure S1 – Research Protocol

2.	Structural Neuroimaging Changes

- Figures S2-4

- Table S1

3.	Morphologic and Cellular Changes

- Figure S5-7

- Tables S2, 3

4.	Integration of Structural Neuroimaging and mRNA Brain Expression Maps

- Tables S4,5

5. Behavioural Changes

-Table S6

6. Weight Changes

- Tables S7,8

*References*

### 1. Materials and Methods

#### Animals: For this study two mouse lines were used, C57BL/6-Tyr^c-Brd^-Tg(Tlr2-luc/gfp)^Kri^/Gaj and C57BL/6-Tlr2^tm1Kir^, named in the text as TLR2 and TLR2^-/-^ respectively. C57BL/6-Tyr^c-Brd^-Tg(Tlr2-luc/gfp)^Kri^/Gaj mouse line was generated in two steps. Firstly, the transgenic mouse model was generated using a Tlr2 promotor and bicistronic luc/gfp transporter on a C57BL/6 genetic background, as previously described.(1) The line was backcrossed to a C57BL/6-Tyr^c-Brd^ mouse background with the goal of enhancing the capture and precision of analysis of the bioluminescent signal in the albino C57BL/6 mice which have a mutation in the tyrosinase gene that enables the photons to penetrate better through the fur of the animal. The second mouse line used was C57BL/6-Tlr2^tm1Kir^, a C57BL/6 mouse line with the Tlr2 gene knocked out on both alleles (Jackson Laboratory, Maine, USA). All the mice used were male, 2-4 months old, bred at the animal facility at the Croatian Institute for Brain Research. The day/ night cycle was defined as 12h -12h, from 7 a.m. to 7 p.m. Food and water were available ad libitum. The animals were housed in transparent polysulphane cages of European standard type 3 (1290D, Tecniplast, Italy). Ten mice were housed in each cage, with the bedding changed every two days.

#### Experimental Design: All animal experiments were done with the permission of the Ethics committee of the University of Zagreb, School of Medicine, ethical permit no. 380-59-10106-14-55/230.

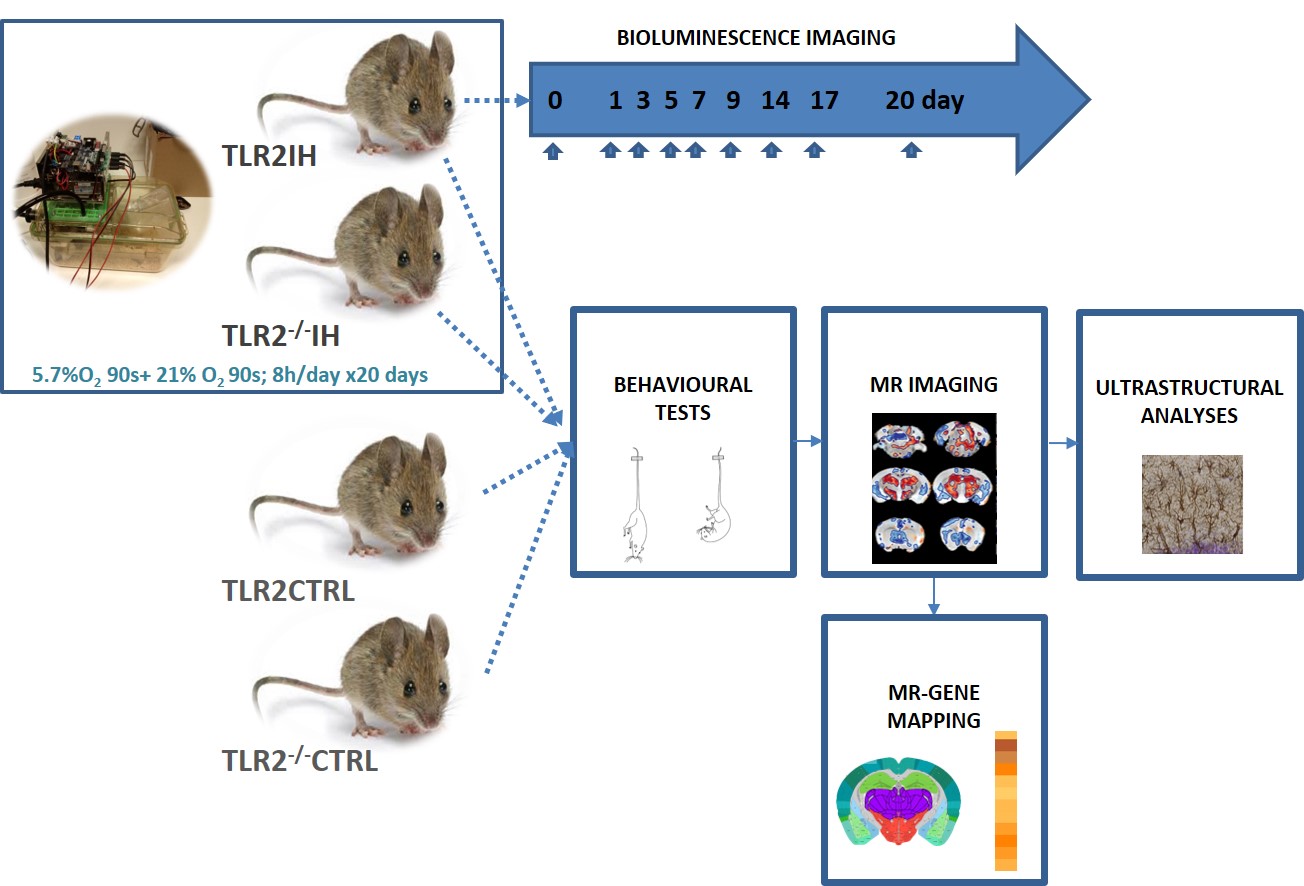

##### Figure S1 Schematic presentation of the research protocol. Four experimental groups were compared in this study: TLR2IH (C57BL/6-Tyr^c-2J^(Cg)-Tg(Tlr2-luc/gfp) 275S Kri), mice with functional TLR2 system that were exposed to three weeks of chronic IH protocol; TLR2CTRL (also C57BL/6-Tyr^c-2J^(Cg)-Tg(Tlr2-luc/gfp) 275S Kri), mice with functional TLR2 system that were handled under control (CTRL) conditions; TLR2^-/-^ IH (C57BL/6-Tlr2^tm1Kir^), TLR2 knock out mice exposed to three weeks of chronic IH; and TLR2^-/-^ CTRL (also C57BL/6-Tlr2^tm1Kir^) control TLR2 knock out mice.

#### Perfusion and Histology For MRI: The mice were killed by transcardiac perfusion with ice-cold heparinized saline followed by 4% buffered paraformaldehyde (PFA). The heads were removed, stored in paraformaldehyde (PFA) for 24h and then rehydrated in Phosphate Buffered Saline (PBS) with 0.05% sodium azide at -4ºC for a minimum of 30 days(2) .

***Behavioural tests:*** *Open Field test* (OF): To determine the spontaneous horizontal locomotor activity, an open-field test was performed, and all the test parameters calculated, as previously described(3). The test was carried out in clear black Plexiglas boxes (40 × 40 × 40 cm) equipped with the video-based system. The test box was cleaned with 70% ethanol between each test. *Y-maze test* (YM): To assess the working memory, a Y-maze test was conducted and all the test parameters calculated, as previously described(3). Arms were cleaned with 70% ethanol between each test to remove odours and residues. The alternation score (%) for each mouse was defined as the ratio of the actual number of alternations to the possible number (defined as the total number of arm entries minus two) multiplied by 100 as shown by the following equation: % Alternation = (Number of alternations)/(Total arm entries - 2) × 100. The number of arm entries was used as an indicator of locomotor activity.(3) *Tail Suspension test* (TST): The TST was performed to evaluate depression-like behaviour. We recorded the overall time that animals were immobile while suspended by the tail over a 6 min period. Scoring of immobility time was performed by means of automated video tracking software as described previously.(4)

#### MRI acquistion: MR images were acquired on a 7 Tesla scanner (Agilent). Samples were immersed in fluorinated liquid to reduce susceptibility artefacts (Galden; Solvay) and loaded four at a time into a 39 mm diameter transmit-receive birdcage coil (Rapid GmbH). High resolution quantitative T1 and T2 maps were acquired using a modified DESPOT1 and DESPOT2-FM protocol(5). This consisted of Spoiled Gradient Recalled (SPGR) images with TE/TR=14.6/32 ms, readout bandwidth 10kHz and seven flip-angles (5-35 degrees in 5 degree steps) and balanced Steady-State Free Precession (bSSFP) images with TE/TR = 4/8 ms, readout bandwidth 62.5 kHz, seven flip angles (8,12,16,24,32,40 & 48 degrees) and four phase increments (45,135,225 and 315 degrees). The flip-angles were chosen to lie between the optimum values for the expected values of T1&T2(6) , which were obtained from a similar preparatory scan. Both SPGR and SSFP had 256x256x256 matrix size with 125 micron isotropic voxels. An Actual Flip-angle Imaging (AFI) scan was acquired for B1 inhomogeneity correction at matrix size 96x96x96, 333 micron isotropic voxel sizes, TE/TR1/TR2 = 6.52/20/100 ms, readout bandwidth 10 kHz and flip-angle 55 degrees(7, 8)

***MRI analysis:*** The MR images were first converted to NIFTI format and then processed using a combination of FSL(9), ANTs(10) and the QUIT toolbox(11). The processing pipeline consisted of several steps, described in detail previously by our group(12). Briefly, images were Tukey filtered in k-space, and then B1, T1 & T2 maps calculated from the AFI, SPGR and SSFP scans respectively. Finally, a synthetic Spin Echo image was calculated from the T1 & T2 maps with TE/TR = 40/10000 ms for registration purposes. This created an almost purely T2-weighted image that matched the contrast of a widely available atlas image(13).The combined 4 images were split into individual subjects and rigidly registered to the atlas to ensure approximate alignment. A template image was then constructed from all subjects in the study(10), which was non-linearly registered to the atlas image. Logarithmic Jacobian determinants were calculated from the inverse warp fields in standard space to estimate apparent volume change. The combined transforms from native to atlas space were applied to all T1 and T2 maps, which were then smoothed with a Gaussian (150 micron FWHM). A brain parenchyma mask was created from the atlas labels by excluding cerebrospinal fluid (CSF) regions. The inverse of the combined transforms for each subject was applied to the atlas mask, and the resulting subject-specific masks were used to calculate the brain and ROIs volume for each subject.

#### A group analysis was then carried out on Jacobian determinant images with permutation tests and Threshold-Free Cluster Enhancement (TFCE) using FSL randomize(14, 15) . The brain volume estimates were included as a regressor of no interest in the design matrix when analysing the Jacobian determinants.

Data are displayed on the mouse template image, using the dual coding approach(16): differences are mapped to color hue, and associated t statistics are mapped to color transparency. Contours are family wise error (FWE) corrected statistically (*P*<.05) significant differences.

#### Neuroplastin immunoreactivity: Coronal sections of 4 animals per each group (four groups were: 1) TLR2 CTRL, 2) TLR2 IH, 3) TLR^-/-^ CTRL and 4) TLR^-/-^ IH) were incubated with 1% hydrogen peroxide in PBS for 30 minutes. After washing, section were incubated in blocking solution (5% horse serum in PBS) for 2 hours at +4°C. Incubation with primary anti‐neuroplastin 65 antibody raised in goat (1:500, R&D Systems, AF5360, Minneapolis, Minnesota, USA) in blocking solution was performed at +4°C overnight. Parallel sections incubated in blocking solution without primary antibody were used as negative controls. Incubation in secondary anti‐goat antibody conjugated with horse‐radish peroxidase (1:10 000, Jackson ImmunoResearch Laboratories, West Grove, Pennsylvania, USA) in blocking solution was performed at RT for 2 hours. Diaminobenzidine (DAB) was used as an enhancement agent for immunoreactivity visualization. Signal was intensified with additional incubation with 0.4% CuSO4. Sections were scanned using a high resolution scanner (Hamamatsu NanoZoomer C10730‐12). Signal intensities of Np immunoreactivity were quantified using ImageJ densitometric analysis (ImageJ, NIH public domain, ttps://imagej.nih.gov/ij/). For the image calibration, a calibrated optical density step tablet was used according to instructions. Statistical analysis of total Np immunoreactivity was done by using the Student's t test. All statistics were performed by using IBM SPSS ver. 25 (IBM Analytics, New York, NY, USA) software.

#### Statistical Analyses: The Kolmogorov–Smirnov test was used to test the normality of distributions. To analyze differences in a variety clinical parameters between investigated groups, one-way analysis of variance with additional post hoc range tests and pairwise multiple comparisons was used to determine which means differ between groups. Bonferroni correction was used to perform pairwise comparisons between group means and to control overall error rate by setting the error rate for each test to the experimentwise error rate divided by the total number of tests. Hence, the observed significance level is adjusted for the fact that multiple comparisons are being made. Pearson correlation coefficients were calculated beween all investigated variables and used to detrmine heatmaps. All statistical analyses had a 2-tailed α level of < .05 for defining significance and were performed by an experienced biostatistician(M.M.) on the statistical software IBM SPSS Statistics version 23 ([www.spss.com](http://www.spss.com" \t "_blank)).

#### 2. Structural Neuroimaging Changes

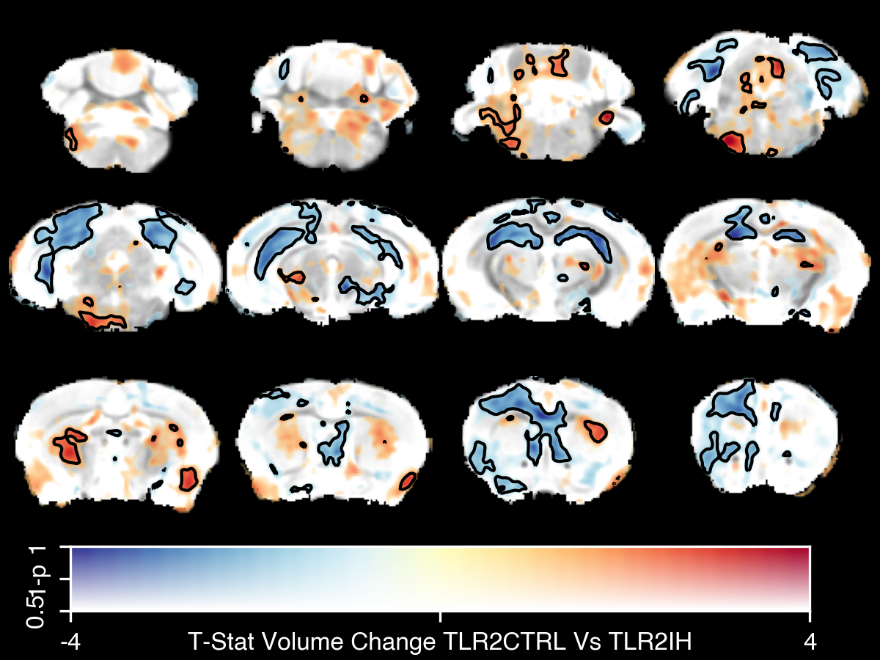

**A**

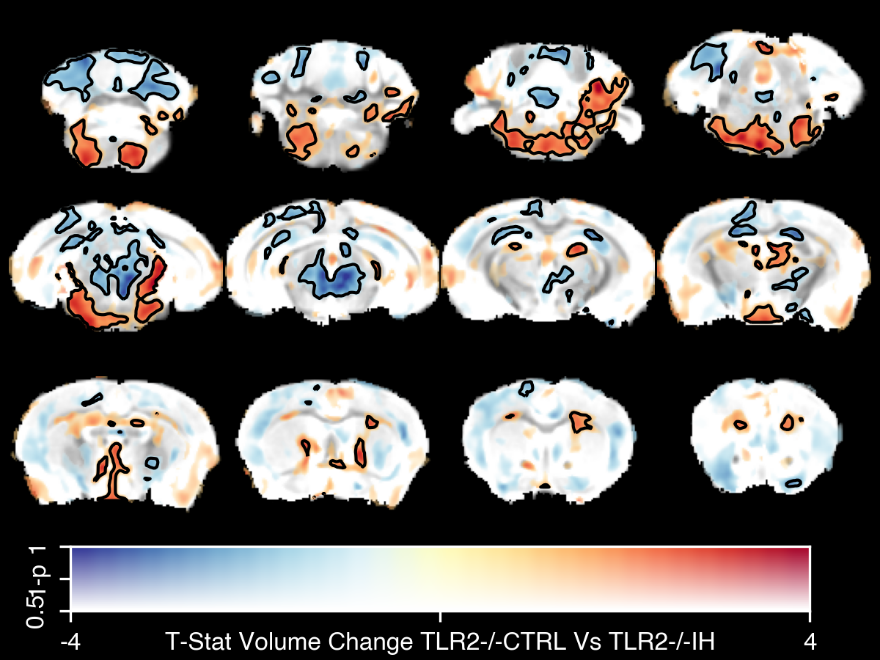

**B**

##### Figure S2 Differences in local brain volume between IH and CTRL conditions in TRL2 (A) and TLR2 ^-/-^ (B) mice. Data are shown as dual-coded statistical maps in which effect size is coded by color hue, and the family-wise error (FWE) corrected p-values are coded by transparency. Cold (blue) colors indicate larger volume in IH mice. Areas in which FWE-corrected P < .05 are contoured in black. Non-parametric statistics were performed using FSL randomize with 5000 permutations and threshold-free cluster enhancement. Hippocampus, left side of the motor and cingulate cortex and septum all appeared enlarged following IH, while these changes are not visible in TLR2 ^-/-^ mice. Reduction of volume after IH is visible in the reticular nuclei of the thalamus and dorsal striatum of TRL2, and pronounced in pons, medulla and mesencephalon of TLR2 ^-/-^.

##### Table S1 Volumetric values of ROIs in grey and white matter in TLR2IH (1; n=14), TLRCTRL (2; n=11), TLR2^-/-^ IH (3; n=15), TLR2^-/-^ CTRL (4; n=8). Values are given as percentage of total brain volume.

*Abbreviations***: Amy-** amygdala; **AC ant-** anterior commissure: pars anterior; **AC post-** anterior commissure: pars posterior; **AV-** arbor vita of cerebellum; **Bsl FB-** basal forebrain; **BNST-** bed nucleus of stria terminalis; **Cbl Ctx-** cerebellar cortex; **Inf Cb Ped-** cerebellar peduncle: inferior; **M Cb Ped-** cerebellar peduncle: middle; **Sup Cb Ped-** cerebellar peduncle: superior; **Cbr Aq-** cerebral aqueduct; **Enr Cx-** cerebral cortex: entorhinal cortex; **F Cx-** cerebral cortex: frontal lobe; **Occ Cx-** cerebral cortex: occipital lobe; **Ptemp Cx-** cerebral cortex: parieto-temporal lobe; **Cbr Ped-** cerebral peduncle; **Inf coll-** colliculus: inferior; **Sup Coll-** colliculus: superior; **CC-** corpus callosum; **CST-** corticospinal tract/pyramids; **Cuneate N-** cuneate nucleus; **DG-** dentate gyrus of hippocampus; **Fac N-** facial nerve (cranial nerve 7); **Fsc Rflx-** fasciculus retroflexus; **Fmb-** fimbria; **Fx-** fornix; **4^th^ Vtr-** fourth ventricle; **F Str-** fundus of striatum; **GP-** globus pallidus; **Hab-** habenular commissure; **Hippo-** hippocampus; **Hypo-** hypothalamus; **Inf Oli-** inferior olivary complex; **Int Cap-** internal capsule; **IntPed Nuc-** interpedunclar nucleus; **Lat Olf T-** lateral olfactory tract; **Lat Spt-** lateral septum; **Lat Vtr-** lateral ventricle; **Mamm-** mammillary bodies; **Mamm Tr-** mammilothalamic tract; **ML-** medial lemniscus/medial longitudinal fasciculus; **Med Spt-** medial septum; **Medulla-** medulla; **Midbrain-** midbrain; **Nac-** nucleus accumbens; **Olf Blb-** olfactory bulbs; **Olf Tub-** olfactory tubercle; **Opt Tr-** optic tract; **Pag-** periaqueductal grey; **Pons-** pons; **Pon N-** pontine nucleus; **Post Com-** posterior commissure; **Pre ParaSub-** pre-para subiculum; **SG Hippo-** stratum granulosum of hippocampus; **Stria M-** stria medullaris; **Stria T-** stria terminalis; **Str-** striatum; **Sub Ependy-** subependymale zone / rhinocele; **Sup Oli-** superior olivary complex; **Thal-** thalamus; **3^rd^ Vtr-** third ventricle; **vHippo** -ventral hippocampus (head); **VT dec-** ventral tegmental decussation.

|  | | **Mean** | **SD** |
| --- | --- | --- | --- |
| LACant | 1 | 0.159 | 0.008 |
|  | 2 | 0.154 | 0.008 |
|  | 3 | 0.147 | 0.010 |
|  | 4 | 0.152 | 0.010 |
| LACpost | 1 | 0.054 | 0.005 |
|  | 2 | 0.052 | 0.005 |
|  | 3 | 0.049 | 0.004 |
|  | 4 | 0.050 | 0.004 |
| LAmy | 1 | 1.740 | 0.060 |
|  | 2 | 1.760 | 0.048 |
|  | 3 | 1.655 | 0.071 |
|  | 4 | 1.664 | 0.038 |
| LAV | 1 | 0.991 | 0.036 |
|  | 2 | 0.983 | 0.075 |
|  | 3 | 1.030 | 0.046 |
|  | 4 | 1.034 | 0.034 |
| LBNST | 1 | 0.156 | 0.010 |
|  | 2 | 0.149 | 0.008 |
|  | 3 | 0.152 | 0.008 |
|  | 4 | 0.154 | 0.006 |
| LBsFB | 1 | 0.526 | 0.020 |
|  | 2 | 0.525 | 0.032 |
|  | 3 | 0.532 | 0.025 |
|  | 4 | 0.522 | 0.041 |
| LCbCtx | 1 | 5.198 | 0.111 |
|  | 2 | 5.184 | 0.180 |
|  | 3 | 5.227 | 0.138 |
|  | 4 | 5.224 | 0.240 |
| LCbrPed | 1 | 0.258 | 0.013 |
|  | 2 | 0.254 | 0.020 |
|  | 3 | 0.267 | 0.010 |
|  | 4 | 0.269 | 0.017 |
| LCC | 1 | 1.783 | 0.046 |
|  | 2 | 1.776 | 0.098 |
|  | 3 | 1.787 | 0.056 |
|  | 4 | 1.798 | 0.041 |
| LCST | 1 | 0.175 | 0.009 |
|  | 2 | 0.179 | 0.010 |
|  | 3 | 0.171 | 0.011 |
|  | 4 | 0.181 | 0.013 |
| LCuneateN | 1 | 0.029 | 0.002 |
|  | 2 | 0.032 | 0.003 |
|  | 3 | 0.030 | 0.004 |
|  | 4 | 0.029 | 0.002 |
| LDG | 1 | 0.419 | 0.021 |
|  | 2 | 0.399 | 0.021 |
|  | 3 | 0.437 | 0.028 |
|  | 4 | 0.427 | 0.021 |
| LEnRCx | 1 | 1.151 | 0.056 |
|  | 2 | 1.160 | 0.058 |
|  | 3 | 1.123 | 0.049 |
|  | 4 | 1.124 | 0.040 |
| LFCx | 1 | 4.698 | 0.187 |
|  | 2 | 4.657 | 0.123 |
|  | 3 | 4.574 | 0.175 |
|  | 4 | 4.481 | 0.168 |
| LFacN | 1 | 0.026 | 0.002 |
|  | 2 | 0.026 | 0.001 |
|  | 3 | 0.026 | 0.001 |
|  | 4 | 0.027 | 0.001 |
| LFscRFlx | 1 | 0.028 | 0.002 |
|  | 2 | 0.028 | 0.002 |
|  | 3 | 0.029 | 0.002 |
|  | 4 | 0.029 | 0.001 |
| LFmb | 1 | 0.303 | 0.015 |
|  | 2 | 0.311 | 0.016 |
|  | 3 | 0.320 | 0.018 |
|  | 4 | 0.322 | 0.016 |
| LFStr | 1 | 0.025 | 0.002 |
|  | 2 | 0.023 | 0.003 |
|  | 3 | 0.024 | 0.003 |
|  | 4 | 0.022 | 0.002 |
| LFx | 1 | 0.066 | 0.004 |
|  | 2 | 0.066 | 0.005 |
|  | 3 | 0.069 | 0.004 |
|  | 4 | 0.069 | 0.003 |
| LGP | 1 | 0.324 | 0.019 |
|  | 2 | 0.324 | 0.012 |
|  | 3 | 0.338 | 0.015 |
|  | 4 | 0.339 | 0.016 |
| LHab | 1 | 0.003 | 0.001 |
|  | 2 | 0.003 | 0.001 |
|  | 3 | 0.003 | 0.001 |
|  | 4 | 0.003 | 0.000 |
| LHippo | 1 | 2.251 | 0.085 |
|  | 2 | 2.154 | 0.125 |
|  | 3 | 2.242 | 0.077 |
|  | 4 | 2.219 | 0.083 |
| LvHippo | 1 | 0.2761 | 0.0186 |
|  | 2 | 0.2639 | 0.0140 |
|  | 3 | 0.2939 | 0.0134 |
|  | 4 | 0.2789 | 0.0191 |
| LHypo | 1 | 1.183 | 0.049 |
|  | 2 | 1.175 | 0.042 |
|  | 3 | 1.216 | 0.034 |
|  | 4 | 1.229 | 0.041 |
| LInfCbPed | 1 | 0.083 | 0.007 |
|  | 2 | 0.084 | 0.008 |
|  | 3 | 0.084 | 0.006 |
|  | 4 | 0.088 | 0.009 |
| LInfcoll | 1 | 0.614 | 0.021 |
|  | 2 | 0.613 | 0.026 |
|  | 3 | 0.634 | 0.029 |
|  | 4 | 0.622 | 0.017 |
| LInfOli | 1 | 0.048 | 0.003 |
|  | 2 | 0.046 | 0.004 |
|  | 3 | 0.045 | 0.005 |
|  | 4 | 0.051 | 0.008 |
| LIntC | 1 | 0.294 | 0.018 |
|  | 2 | 0.300 | 0.011 |
|  | 3 | 0.305 | 0.013 |
|  | 4 | 0.305 | 0.016 |
| LLatSpt | 1 | 0.373 | 0.025 |
|  | 2 | 0.345 | 0.027 |
|  | 3 | 0.335 | 0.016 |
|  | 4 | 0.337 | 0.018 |
| LLatVtr | 1 | 0.420 | 0.029 |
|  | 2 | 0.414 | 0.025 |
|  | 3 | 0.405 | 0.021 |
|  | 4 | 0.404 | 0.022 |
| LLatOlfT | 1 | 0.141 | 0.012 |
|  | 2 | 0.147 | 0.010 |
|  | 3 | 0.144 | 0.012 |
|  | 4 | 0.147 | 0.022 |
| LMCbPed | 1 | 0.139 | 0.007 |
|  | 2 | 0.137 | 0.010 |
|  | 3 | 0.141 | 0.007 |
|  | 4 | 0.146 | 0.006 |
| LMamm | 1 | 0.056 | 0.006 |
|  | 2 | 0.057 | 0.005 |
|  | 3 | 0.058 | 0.005 |
|  | 4 | 0.058 | 0.002 |
| LMammTr | 1 | 0.028 | 0.002 |
|  | 2 | 0.027 | 0.002 |
|  | 3 | 0.030 | 0.003 |
|  | 4 | 0.030 | 0.002 |
| LMedSpt | 1 | 0.122 | 0.010 |
|  | 2 | 0.123 | 0.010 |
|  | 3 | 0.119 | 0.007 |
|  | 4 | 0.117 | 0.010 |
| LML | 1 | 0.256 | 0.008 |
|  | 2 | 0.253 | 0.013 |
|  | 3 | 0.261 | 0.011 |
|  | 4 | 0.263 | 0.016 |
| LNac | 1 | 0.422 | 0.015 |
|  | 2 | 0.412 | 0.027 |
|  | 3 | 0.406 | 0.033 |
|  | 4 | 0.396 | 0.012 |
| LOccCx | 1 | 0.622 | 0.029 |
|  | 2 | 0.607 | 0.036 |
|  | 3 | 0.605 | 0.021 |
|  | 4 | 0.598 | 0.013 |
| LOlfBlb | 1 | 2.783 | 0.103 |
|  | 2 | 2.822 | 0.119 |
|  | 3 | 2.789 | 0.087 |
|  | 4 | 2.815 | 0.097 |
| LOptTr | 1 | 0.187 | 0.007 |
|  | 2 | 0.183 | 0.008 |
|  | 3 | 0.186 | 0.008 |
|  | 4 | 0.188 | 0.008 |
| LOlfTub | 1 | 0.388 | 0.025 |
|  | 2 | 0.386 | 0.029 |
|  | 3 | 0.387 | 0.024 |
|  | 4 | 0.384 | 0.032 |
| LPonN | 1 | 0.084 | 0.006 |
|  | 2 | 0.086 | 0.009 |
|  | 3 | 0.089 | 0.009 |
|  | 4 | 0.095 | 0.007 |
| LPreParaSub | 1 | 0.281 | 0.019 |
|  | 2 | 0.259 | 0.022 |
|  | 3 | 0.282 | 0.016 |
|  | 4 | 0.279 | 0.013 |
| LPtempCx | 1 | 8.023 | 0.184 |
|  | 2 | 8.063 | 0.261 |
|  | 3 | 7.879 | 0.211 |
|  | 4 | 7.924 | 0.197 |
| LSGHippo | 1 | 0.109 | 0.007 |
|  | 2 | 0.105 | 0.007 |
|  | 3 | 0.115 | 0.006 |
|  | 4 | 0.115 | 0.007 |
| LStr | 1 | 2.265 | 0.046 |
|  | 2 | 2.310 | 0.046 |
|  | 3 | 2.270 | 0.057 |
|  | 4 | 2.285 | 0.047 |
| LStriaM | 1 | 0.073 | 0.006 |
|  | 2 | 0.077 | 0.004 |
|  | 3 | 0.081 | 0.006 |
|  | 4 | 0.081 | 0.003 |
| LStriaT | 1 | 0.087 | 0.005 |
|  | 2 | 0.090 | 0.005 |
|  | 3 | 0.092 | 0.006 |
|  | 4 | 0.094 | 0.004 |
| LSubEpendy | 1 | 0.008 | 0.001 |
|  | 2 | 0.008 | 0.001 |
|  | 3 | 0.007 | 0.001 |
|  | 4 | 0.008 | 0.000 |
| LSupCbPed | 1 | 0.099 | 0.003 |
|  | 2 | 0.100 | 0.004 |
|  | 3 | 0.103 | 0.005 |
|  | 4 | 0.100 | 0.007 |
| LSupColl | 1 | 0.850 | 0.026 |
|  | 2 | 0.853 | 0.032 |
|  | 3 | 0.924 | 0.039 |
|  | 4 | 0.910 | 0.033 |
| LSupOli | 1 | 0.096 | 0.008 |
|  | 2 | 0.097 | 0.007 |
|  | 3 | 0.095 | 0.007 |
|  | 4 | 0.097 | 0.008 |
| LThal | 1 | 1.786 | 0.069 |
|  | 2 | 1.829 | 0.070 |
|  | 3 | 1.891 | 0.059 |
|  | 4 | 1.917 | 0.041 |
| RACant | 1 | 0.127 | 0.008 |
|  | 2 | 0.127 | 0.005 |
|  | 3 | 0.124 | 0.007 |
|  | 4 | 0.122 | 0.007 |
| RACpost | 1 | 0.053 | 0.004 |
|  | 2 | 0.050 | 0.002 |
|  | 3 | 0.051 | 0.006 |
|  | 4 | 0.052 | 0.003 |
| RAmy | 1 | 1.660 | 0.064 |
|  | 2 | 1.704 | 0.110 |
|  | 3 | 1.640 | 0.059 |
|  | 4 | 1.631 | 0.100 |
| RAV | 1 | 1.041 | 0.033 |
|  | 2 | 1.038 | 0.032 |
|  | 3 | 1.064 | 0.065 |
|  | 4 | 1.031 | 0.039 |
| RBNST | 1 | 0.142 | 0.006 |
|  | 2 | 0.137 | 0.006 |
|  | 3 | 0.136 | 0.008 |
|  | 4 | 0.140 | 0.005 |
| RBsFB | 1 | 0.518 | 0.023 |
|  | 2 | 0.506 | 0.016 |
|  | 3 | 0.533 | 0.026 |
|  | 4 | 0.526 | 0.032 |
| RCbCtx | 1 | 5.451 | 0.177 |
|  | 2 | 5.424 | 0.182 |
|  | 3 | 5.392 | 0.168 |
|  | 4 | 5.245 | 0.088 |
| RCbrPed | 1 | 0.233 | 0.018 |
|  | 2 | 0.241 | 0.013 |
|  | 3 | 0.250 | 0.011 |
|  | 4 | 0.245 | 0.010 |
| RCC | 1 | 1.919 | 0.072 |
|  | 2 | 1.886 | 0.067 |
|  | 3 | 1.926 | 0.049 |
|  | 4 | 1.923 | 0.063 |
| RCST | 1 | 0.173 | 0.009 |
|  | 2 | 0.175 | 0.009 |
|  | 3 | 0.176 | 0.013 |
|  | 4 | 0.179 | 0.015 |
| RCuneateN | 1 | 0.027 | 0.002 |
|  | 2 | 0.029 | 0.002 |
|  | 3 | 0.030 | 0.004 |
|  | 4 | 0.029 | 0.002 |
| RDG | 1 | 0.401 | 0.023 |
|  | 2 | 0.377 | 0.018 |
|  | 3 | 0.406 | 0.015 |
|  | 4 | 0.397 | 0.025 |
| REnRCx | 1 | 1.318 | 0.042 |
|  | 2 | 1.304 | 0.056 |
|  | 3 | 1.256 | 0.049 |
|  | 4 | 1.268 | 0.066 |
| RFCx | 1 | 5.061 | 0.113 |
|  | 2 | 4.956 | 0.266 |
|  | 3 | 4.859 | 0.130 |
|  | 4 | 4.796 | 0.142 |
| RFacN | 1 | 0.023 | 0.001 |
|  | 2 | 0.023 | 0.001 |
|  | 3 | 0.023 | 0.001 |
|  | 4 | 0.024 | 0.002 |
| RFscRFlx | 1 | 0.030 | 0.002 |
|  | 2 | 0.031 | 0.003 |
|  | 3 | 0.031 | 0.002 |
|  | 4 | 0.030 | 0.001 |
| RFmb | 1 | 0.326 | 0.016 |
|  | 2 | 0.339 | 0.026 |
|  | 3 | 0.350 | 0.020 |
|  | 4 | 0.363 | 0.019 |
| RFStr | 1 | 0.021 | 0.002 |
|  | 2 | 0.020 | 0.003 |
|  | 3 | 0.021 | 0.003 |
|  | 4 | 0.020 | 0.002 |
| RFx | 1 | 0.069 | 0.004 |
|  | 2 | 0.070 | 0.003 |
|  | 3 | 0.071 | 0.004 |
|  | 4 | 0.072 | 0.005 |
| RGP | 1 | 0.321 | 0.026 |
|  | 2 | 0.340 | 0.019 |
|  | 3 | 0.354 | 0.017 |
|  | 4 | 0.355 | 0.021 |
| RHab | 1 | 0.004 | 0.001 |
|  | 2 | 0.004 | 0.001 |
|  | 3 | 0.004 | 0.001 |
|  | 4 | 0.004 | 0.000 |
| RHippo | 1 | 2.316 | 0.061 |
|  | 2 | 2.231 | 0.076 |
|  | 3 | 2.305 | 0.062 |
|  | 4 | 2.290 | 0.102 |
| RvHippo | 1 | 0.2814 | 0.0124 |
|  | 2 | 0.2728 | 0.0134 |
|  | 3 | 0.3108 | 0.0157 |
|  | 4 | 0.3081 | 0.0190 |
| RHypo | 1 | 1.110 | 0.049 |
|  | 2 | 1.111 | 0.043 |
|  | 3 | 1.149 | 0.033 |
|  | 4 | 1.173 | 0.043 |
| RInfCbPed | 1 | 0.081 | 0.007 |
|  | 2 | 0.084 | 0.007 |
|  | 3 | 0.081 | 0.008 |
|  | 4 | 0.085 | 0.007 |
| RInfcoll | 1 | 0.649 | 0.019 |
|  | 2 | 0.651 | 0.020 |
|  | 3 | 0.659 | 0.027 |
|  | 4 | 0.643 | 0.019 |
| RInfOli | 1 | 0.042 | 0.005 |
|  | 2 | 0.042 | 0.003 |
|  | 3 | 0.042 | 0.003 |
|  | 4 | 0.044 | 0.006 |
| RIntC | 1 | 0.257 | 0.013 |
|  | 2 | 0.275 | 0.010 |
|  | 3 | 0.278 | 0.010 |
|  | 4 | 0.285 | 0.018 |
| RLatSpt | 1 | 0.357 | 0.022 |
|  | 2 | 0.342 | 0.017 |
|  | 3 | 0.335 | 0.019 |
|  | 4 | 0.331 | 0.016 |
| RLatVtr | 1 | 0.364 | 0.019 |
|  | 2 | 0.377 | 0.026 |
|  | 3 | 0.355 | 0.020 |
|  | 4 | 0.373 | 0.027 |
| RLatOlfT | 1 | 0.143 | 0.008 |
|  | 2 | 0.149 | 0.010 |
|  | 3 | 0.144 | 0.012 |
|  | 4 | 0.143 | 0.010 |
| RMCbPed | 1 | 0.128 | 0.006 |
|  | 2 | 0.133 | 0.003 |
|  | 3 | 0.130 | 0.006 |
|  | 4 | 0.132 | 0.004 |
| RMamm | 1 | 0.068 | 0.005 |
|  | 2 | 0.072 | 0.006 |
|  | 3 | 0.072 | 0.004 |
|  | 4 | 0.071 | 0.005 |
| RMammTr | 1 | 0.026 | 0.002 |
|  | 2 | 0.026 | 0.002 |
|  | 3 | 0.027 | 0.002 |
|  | 4 | 0.027 | 0.002 |
| RMedSpt | 1 | 0.141 | 0.011 |
|  | 2 | 0.142 | 0.009 |
|  | 3 | 0.137 | 0.011 |
|  | 4 | 0.131 | 0.010 |
| RML | 1 | 0.267 | 0.012 |
|  | 2 | 0.269 | 0.011 |
|  | 3 | 0.275 | 0.014 |
|  | 4 | 0.272 | 0.011 |
| RNac | 1 | 0.438 | 0.022 |
|  | 2 | 0.422 | 0.018 |
|  | 3 | 0.417 | 0.026 |
|  | 4 | 0.412 | 0.018 |
| ROccCx | 1 | 0.770 | 0.038 |
|  | 2 | 0.766 | 0.029 |
|  | 3 | 0.768 | 0.039 |
|  | 4 | 0.747 | 0.032 |
| ROlfBlb | 1 | 2.896 | 0.124 |
|  | 2 | 2.932 | 0.134 |
|  | 3 | 2.857 | 0.098 |
|  | 4 | 2.935 | 0.123 |
| ROptTr | 1 | 0.168 | 0.009 |
|  | 2 | 0.172 | 0.008 |
|  | 3 | 0.177 | 0.009 |
|  | 4 | 0.182 | 0.005 |
| ROlfTub | 1 | 0.403 | 0.021 |
|  | 2 | 0.402 | 0.023 |
|  | 3 | 0.408 | 0.031 |
|  | 4 | 0.386 | 0.026 |
| RPonN | 1 | 0.083 | 0.006 |
|  | 2 | 0.086 | 0.007 |
|  | 3 | 0.088 | 0.008 |
|  | 4 | 0.089 | 0.005 |
| RPreParaSub | 1 | 0.246 | 0.010 |
|  | 2 | 0.225 | 0.017 |
|  | 3 | 0.245 | 0.012 |
|  | 4 | 0.239 | 0.011 |
| RPtempCx | 1 | 9.106 | 0.287 |
|  | 2 | 9.089 | 0.291 |
|  | 3 | 8.981 | 0.158 |
|  | 4 | 8.852 | 0.247 |
| RSGHippo | 1 | 0.092 | 0.007 |
|  | 2 | 0.088 | 0.006 |
|  | 3 | 0.094 | 0.004 |
|  | 4 | 0.094 | 0.006 |
| RStr | 1 | 2.134 | 0.093 |
|  | 2 | 2.168 | 0.087 |
|  | 3 | 2.148 | 0.066 |
|  | 4 | 2.150 | 0.099 |
| RStriaM | 1 | 0.067 | 0.004 |
|  | 2 | 0.070 | 0.004 |
|  | 3 | 0.073 | 0.003 |
|  | 4 | 0.070 | 0.002 |
| RStriaT | 1 | 0.100 | 0.006 |
|  | 2 | 0.105 | 0.007 |
|  | 3 | 0.112 | 0.008 |
|  | 4 | 0.115 | 0.009 |
| RSubEpendy | 1 | 0.006 | 0.001 |
|  | 2 | 0.006 | 0.001 |
|  | 3 | 0.006 | 0.001 |
|  | 4 | 0.006 | 0.001 |
| RSupCbPed | 1 | 0.110 | 0.004 |
|  | 2 | 0.113 | 0.005 |
|  | 3 | 0.118 | 0.010 |
|  | 4 | 0.112 | 0.006 |
| RSupColl | 1 | 0.815 | 0.031 |
|  | 2 | 0.820 | 0.039 |
|  | 3 | 0.873 | 0.036 |
|  | 4 | 0.858 | 0.037 |
| RSupOli | 1 | 0.088 | 0.005 |
|  | 2 | 0.097 | 0.007 |
|  | 3 | 0.091 | 0.009 |
|  | 4 | 0.104 | 0.008 |
| RThal | 1 | 1.725 | 0.051 |
|  | 2 | 1.743 | 0.070 |
|  | 3 | 1.804 | 0.044 |
|  | 4 | 1.835 | 0.057 |

#### Detailed structural image analysis

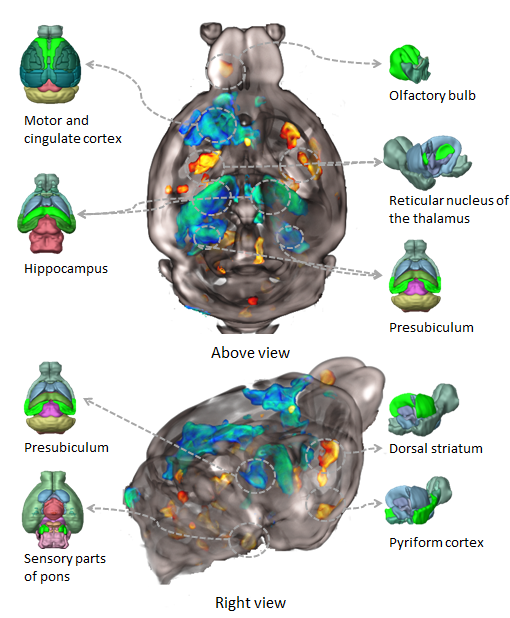

##### Figure S3 Details of structural changes after exposure to chronic intermittent hypoxia in mice with a functional TLR2 gene.

Image shows statistically significant differences between T*LR2 CTRL and TLR2 IH with anatomical models from Brain Explorer 2 for* Allen Mouse Brain Atlas (*http://mouse.brain-map.org/static/atlas). Most significant enlargements are visible bilaterally in the hippocampi and presubiculi and left motor and cingulate cortices while the reduction of volume is evident bilaterally in the reticular nuclei of the thalamus, right dorsal striatum, right piriform cortex and dorsolateral (sensory) parts of the pons (pontine tegmentum).*

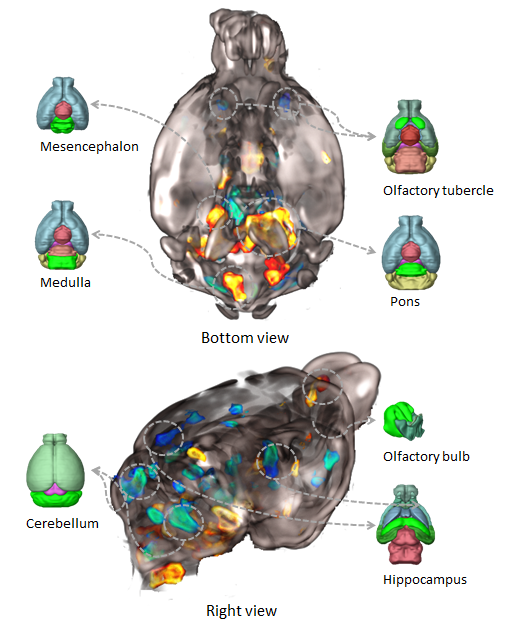

##### Figure S4 Details of structural changes after exposure to chronic intermittent hypoxia in mice without a functional TLR2 gene.

*Image shows statistically significant differences between TLR2^-/-^ CTRL and* *TLR2^-/-^ IH with anatomical models from Brain Explorer 2 for* Allen Mouse Brain Atlas (*http://mouse.brain-map.org/static/atlas). Slight increases are visible in the right hippocampus and presubiculum and both olfactory tubercles. More significant enlargements are seen in the parts of the cerebellum while reductions of volume were noted in the pons, mesencephalon and medulla.*

#### 3. Morphologic and Cellular changes

In our experimental set up, we were unable to demonstrate quantitively gross cellular differences between differential TLR2 genotypes and experimental IH phenotypes to directly account for the primary neuroimaging findings in our study. More specifically, we did not detect gross astroglial proliferation (Figures S4, S5), although a clear protective effect of TLR2 genotype against demyelinating effect of IH experimental protocol was shown (Figure S6).

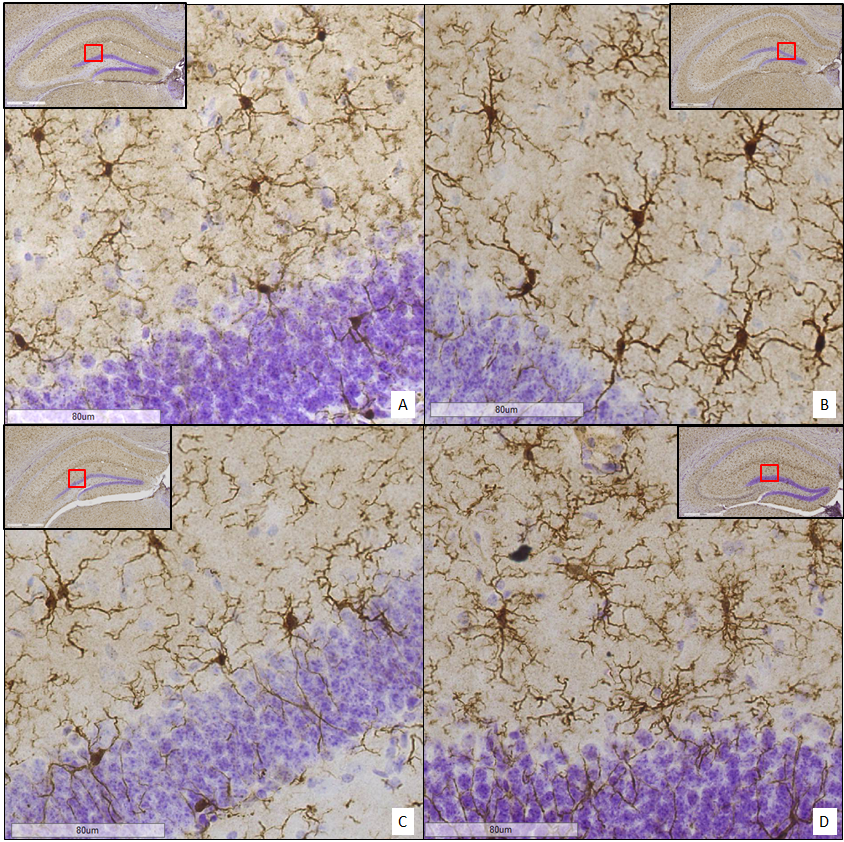

##### Figure S5 Representative images of Iba1 positive cells in the dentate gyrus of four investigated groups.

In the corners of the images are respective areas in red rectangles that the photomicrograph was taken from. Phenotypical distinctions between microglia in four investigated groups are observed, albeit quantatively statistically not found to be significant. A: TLR2IH; B: TLR2 CTRL; C: TLR2^-/-^ IH; D: TLR2^-/-^ CTRL; scale bar denotes 80 µm.

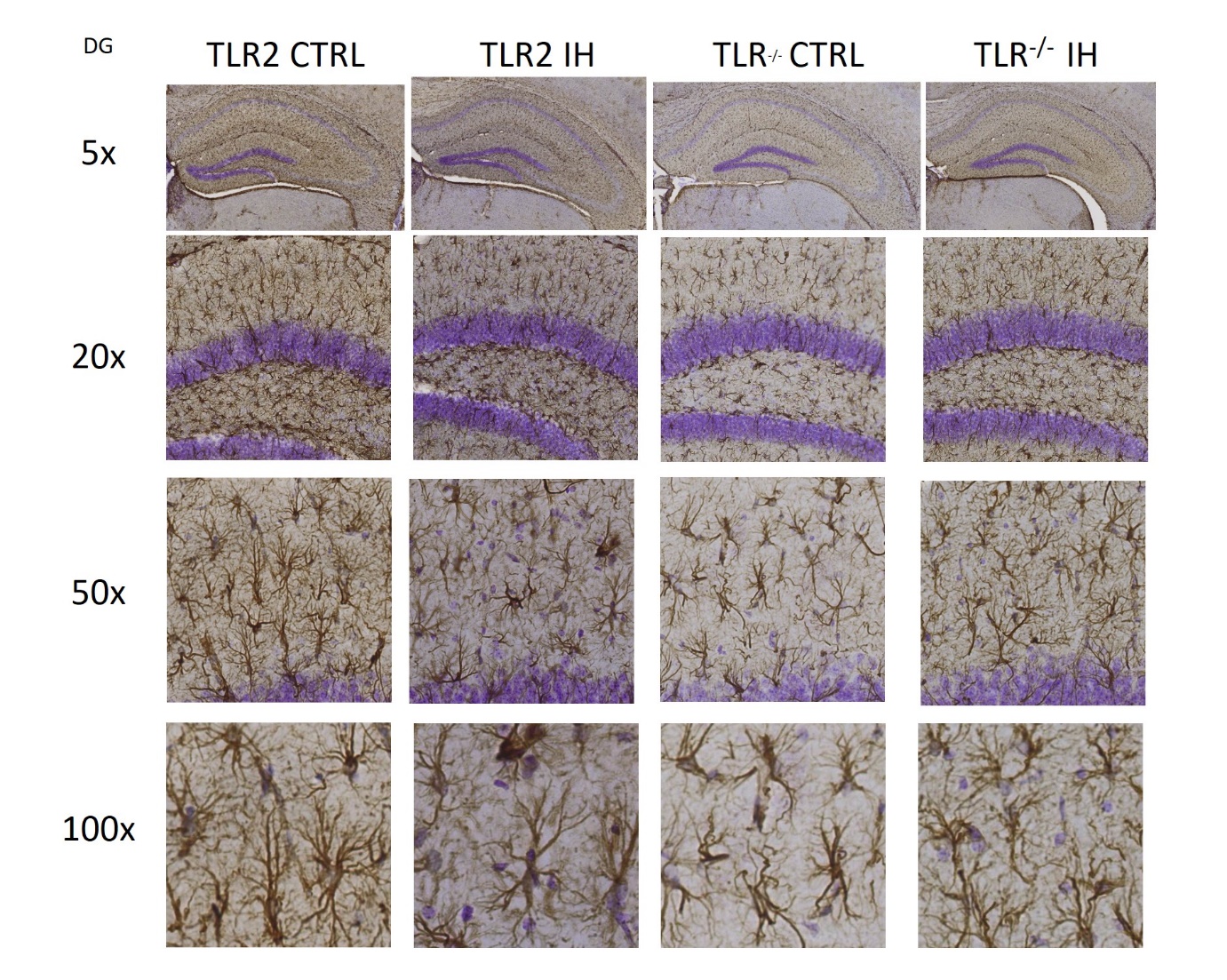

##### Figure S6 Representative images of GFAP positive cells, astrocytes, in the dentate gyrus of hippocampi of four investigated groups under different magnifications.

Phenotypical distinctions between astrocytes in four investigated groups are seen, presumably driven by genotype and genotype-driven modulation of response to IH, albeit quantatively statistically not found to be significant. A: TLR2 IH; B: TLR2 CTRL; C: TLR2^-/-^ IH; D: TLR2^-/-^ CTRL.

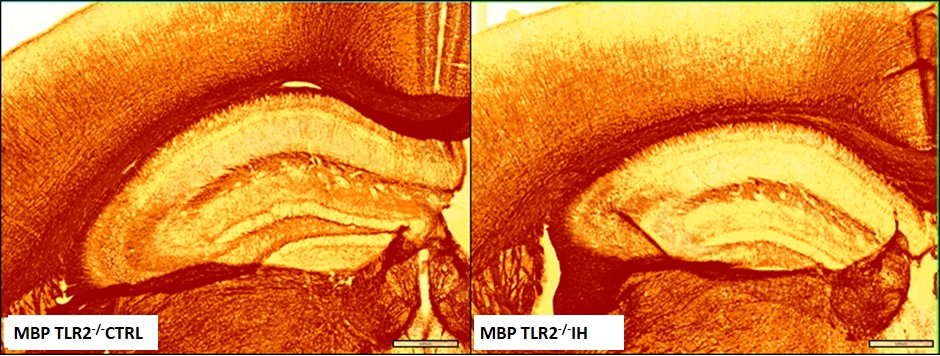

##### Figure S7 Representative images of myelin basic protein staining in the hippocampus. Digital thresholding analysis showed the area of MBP positive area to be significantly higher in the TLR2^-/-^ CTRL group (n (mice)=4; 6 sections) than in the TLR2^-/-^ IH group (n=4;6 sections; P=.0286). (5x; Aperio Image software; scale bar denotes 500 µm)

*Abbreviations*: MBP- myelin basic protein; IH- intermittent hypoxia; CTRL- control; TLR2^-/-^ - TLR2 knock out mouse.

##### Table S2 Quantitative evaluation of several cellular markers in TLR2 IH (1), TLR CTRL (2), TLR2^-/-^ IH (3), TLR2^-/-^ CTRL (4). The values were normalized as per described protocol.

*Abbreviations*: cFos- nuclear phosphoprotein ; CA-cornu ammonis; DG – dentate gyrus; GFAP- Glial fibrillary acidic protein ; MBP- myelin basic protein; mean -mean number of cells; Iba1- ionized calcium binding adaptor molecule 1; IH- intermittent hypoxia; CTRL- control; ROI- region of interest; SD – standard deviation; SEM- standard error of mean; TLR2^-/-^ - TLR2 knock out mouse.

| **Cellular marker: ROI** | |  |  |
| --- | --- | --- | --- |
|  |  | **Mean** | **SD** |
| cFos: hippocampus | 1 | 141848.32 | 36115.70 |
|  | 2 | 146464.27 | 27391.78 |
|  | 3 | 98875.72 | 29380.85 |
|  | 4 | 99912.39 | 38871.37 |
| cFos: hypothalamus | 1 | 61836.00 | 7403.99 |
|  | 2 | 58740.55 | 8032.98 |
|  | 3 | 58813.34 | 7003.78 |
|  | 4 | 69054.76 | 25471.26 |
| cFos: thalamus | 1 | 58890.02 | 4700.41 |
|  | 2 | 56962.43 | 4935.93 |
|  | 3 | 54841.82 | 5283.58 |
|  | 4 | 50594.74 | 8370.90 |
| GFAP: hippocampus | 1 | 25.79 | 6.54 |
|  | 2 | 21.50 | 0.76 |
|  | 3 | 23.26 | 11.82 |
|  | 4 | 29.61 | 11.01 |
| GFAP: hypothalamus | 1 | 15.70 | 6.67 |
|  | 2 | 14.15 | 3.64 |
|  | 3 | 14.01 | 9.85 |
|  | 4 | 17.89 | 7.37 |
| GFAP: thalamus | 1 | 3.74 | 2.15 |
|  | 2 | 3.10 | 1.17 |
|  | 3 | 3.09 | 1.19 |
|  | 4 | 5.55 | 2.82 |
| Iba1: thalamus | 1 | 4415.93 | 704.33 |
|  | 2 | 4612.50 | 614.20 |
|  | 3 | 4464.27 | 663.04 |
|  | 4 | 4755.20 | 498.86 |
| Iba1: hypothalamus | 1 | 4306.10 | 1190.58 |
|  | 2 | 4448.58 | 797.97 |
|  | 3 | 4503.18 | 1084.33 |
|  | 4 | 4945.44 | 677.90 |
| Iba1: DG | 1 | 5187.81 | 339.30 |
|  | 2 | 5739.16 | 1111.37 |
|  | 3 | 6094.35 | 783.67 |
|  | 4 | 6046.41 | 471.45 |
| Iba1: CA1 | 1 | 5911.13 | 729.22 |
|  | 2 | 6011.04 | 853.60 |
|  | 3 | 5702.21 | 928.13 |
|  | 4 | 5683.61 | 232.24 |
| Iba1: CA3 | 1 | 5242.53 | 537.55 |
|  | 2 | 5420.07 | 403.26 |
|  | 3 | 5068.17 | 906.29 |
|  | 4 | 5855.42 | 401.24 |
| lba1: dorsal hippocampus | 1 | 5782.44 | 526.51 |
|  | 2 | 5864.36 | 471.21 |
|  | 3 | 5592.27 | 696.95 |
|  | 4 | 5886.82 | 510.65 |
| lba1: ventral hippocampus | 1 | 5058.64 | 511.75 |
|  | 2 | 5389.90 | 537.60 |
|  | 3 | 4986.64 | 651.43 |
|  | 4 | 5085.61 | 310.45 |
| MBP | 1 | 3,45998 | 1,349705 |
|  | 2 | 5,644926 | 2,301069 |
|  | 3 | 5,320819 | 1,030496 |
|  | 4 | 8,049712 | 1,789514 |

##### Table S3. Average neuroplastin immunoreactivity values as integrated optical density (IOD) are shown for all major hippocampal regions.

*Abbreviations*: DG – dentate gyrus. CTRL – control; CA – Ammon’s horn (*Cornu Ammonis*), IH- intermittent hypoxia, SD- standard deviation, str. – stratum, TLR2^-/-^ - TLR2 knock out mouse.

|  | **REGION** | **TLR2 IH** | | **TLR2 CTRL** | | **TLR2^-/-^ IH** | | **TLR2^-/-^ CTRL** | |
| --- | --- | --- | --- | --- | --- | --- | --- | --- | --- |
|  |  | mean | SD | mean | SD | mean | SD | mean | SD |
| DG | *str. granulare* | 1551837.806 | 92460.08074 | 1625417.514 | 280262.4315 | 1473234.185 | 220753.828 | 1188363.033 | 78114.9897 |
|  | *str. moleculare* | 1357669.236 | 55887.72664 | 1421728.306 | 264159.4369 | 1296908.065 | 147027.1163 | 1082671.422 | 49815.32292 |
| CA3 | *str. pyramidale* | 1480109.458 | 117896.6354 | 1564299.403 | 192327.5423 | 1339265.222 | 152809.9737 | 1145629.033 | 167876.599 |
|  | *str. radiatum* | 1426939.75 | 123935.5783 | 1484091.972 | 174778.397 | 1293531.472 | 140770.0314 | 1133350.922 | 133030.3292 |
|  | *str. oriens* | 1417715.167 | 202002.4107 | 1538039.806 | 160727.3422 | 1288390.306 | 156226.1885 | 1128826.322 | 118559.0546 |
| CA2 | *str. pyramidale* | 1450676.014 | 205226.6957 | 1563036.736 | 199066.5075 | 1363181.694 | 140448.6548 | 1112153.056 | 166972.7896 |
|  | *str. radiatum* | 1375802.9 | 109941.6369 | 1470989.83 | 207154.8051 | 1303567.11 | 150504.0415 | 1118247.89 | 144971.1249 |
|  | *str. oriens* | 1419759.792 | 204980.1828 | 1518213.833 | 172862.7509 | 1335268.722 | 162498.5641 | 1101295.156 | 126852.9329 |
| CA1 | *str. pyramidale* | 1634771.259 | 128044.1863 | 1526307.852 | 178344.4649 | 1444880.037 | 233533.1714 | 1233602.500 | 130180.1678 |
|  | *str. radiatum* | 1481651.407 | 139739.1881 | 1419492.648 | 137166.17 | 1353984.713 | 217919.114 | 1175047.389 | 66198.10435 |
|  | *str. oriens* | 1522651.352 | 74172.47761 | 1483661.815 | 109709.5437 | 1366674.028 | 220241.6968 | 1182495.722 | 144191.7691 |

##### 4. Integration of Structural Neuroimaging and mRNA Brain Expression Maps

##### Table S4. Linear regression results are shown. Green field denote statistical significance. Abbreviations: ARHGEF6- Rho Guanine Nucleotide Exchange Factor 6; BDNF- the brain derived neurotrophic factor; CAMK2A- Calcium/calmodulin dependent protein kinase II alpha; CCK- Cholecystokinin; PCP4; MOBP- Myelin Associated Oligodendrocyte Basic Protein; ; NPTN-neuroplastin FN1- Fibronectin 1; HOMER1; RASGRP1- RAS guanyl nucleotide-releasing protein 1; TLR2^-/-^ - TLR2 knock out mouse WT-wild type (functional TLR2).

| **MR Parameter (group)** | **Variable** | **PCP4** | **MOBP** | **FN1** | **ARHGEF6** | **CAMK2A** | **HOMER1** | **CCK** | **BDNF** | **RASGRP1** | **NPTN** |
| --- | --- | --- | --- | --- | --- | --- | --- | --- | --- | --- | --- |
| Volume (WT) | Pearson Correlation | .103 | -.467 | .635* | .214 | .644* | .491 | .418 | .534 | .580* | .532 |
|  | Sig. (2-tailed) | .751 | .126 | .027 | .504 | .024 | .105 | .201 | .074 | .048 | .075 |
|  | N | 12 | 12 | 12 | 12 | 12 | 12 | 11 | 12 | 12 | 12 |
| V (TLR2 ^-/-^) | Pearson Correlation | .220 | -.297 | .162 | .047 | .376 | .315 | .088 | .132 | .404 | .419 |
|  | Sig. (2-tailed) | .492 | .348 | .615 | .884 | .229 | .318 | .796 | .683 | .193 | .175 |
|  | N | 12 | 12 | 12 | 12 | 12 | 12 | 11 | 12 | 12 | 12 |
| T1 (WT) | Pearson Correlation | .194 | -.468 | .682* | .685* | .650* | .489 | .797** | .813** | .613* | .566 |
|  | Sig. (2-tailed) | .546 | .125 | .015 | .014 | .022 | .107 | .003 | .001 | .034 | .055 |
|  | N | 12 | 12 | 12 | 12 | 12 | 12 | 11 | 12 | 12 | 12 |
| T1(TLR2 ^-/-^) | Pearson Correlation | .157 | -.160 | .505 | .196 | .300 | .104 | .557 | .450 | .253 | .165 |
|  | Sig. (2-tailed) | .626 | .618 | .094 | .541 | .343 | .747 | .075 | .142 | .427 | .608 |
|  | N | 12 | 12 | 12 | 12 | 12 | 12 | 11 | 12 | 12 | 12 |
| T2 (WT) | Pearson Correlation | -.454 | .389 | -.215 | -.332 | -.691* | -.469 | -.454 | -.457 | -.700* | -.559 |
|  | Sig. (2-tailed) | .138 | .211 | .501 | .292 | .013 | .124 | .160 | .136 | .011 | .059 |
|  | N | 12 | 12 | 12 | 12 | 12 | 12 | 11 | 12 | 12 | 12 |
| T2 (TLR2^-/-^) | Pearson Correlation | -.515 | -.002 | .016 | .034 | -.549 | -.237 | -.268 | -.297 | -.522 | -.474 |
|  | Sig. (2-tailed) | .087 | .994 | .961 | .918 | .065 | .457 | .425 | .349 | .082 | .120 |
|  | N | 12 | 12 | 12 | 12 | 12 | 12 | 11 | 12 | 12 | 12 |

* refers to statistically significant results (*P* <.05)

##### Table S5. The list of major neuroplasticity genes that were investigated.

| Group | Genes | Relevant References |
| --- | --- | --- |
| Sleep homeostasis (Neuroplasticity) | BDNF, FOS, FOSB, FOSL1, FOSL2, PAX6, AVPR1B, CCK, TLR2, HOMER1, ARC, EGR1, GSK3B, CAMK2A, GAP43, NGF, DISC1, COMT | (17-23) (24) (25) (26) (27) (28) (29) (30) (31, 32) |
| Neuroplasticity | FIGF, NGFR, NRG1, GSK3A, CREB1, CREB3, CRTC1, GPM6A, NCAM1, NCAM2, REST, RGS14, PCP4, ARHGEF6, DRD3, ITM2B, LRRTM2, MDK, RASGRP1, SLITRK5, SSTR4, NPTN, EPHA2, EMP1, FN1, HES5, SOX11, CORO1A, NR2E1, GNAQ, CLK1, USF2, SRF | (33-58) (59-63) |
| Growth/development | PROX1, SOX2, MSX1 | (64-69) |
| Myelin formation | CLDN11, MOBP | (70), (71), (72), (73) |
| Immunoactive factor | CAMP | (74), (75), (76), (77) |
| Angiogenic factor | ANGPT22 | (78), (79),  (80), (81), (82) |

### 5. Behavioural Changes

##### Table S6. Values of behavioural test parameters in four groups of animals: TLR2 IH (1; n=15), TLR CTRL (2; n=12), TLR2^-/-^ IH (3; n=16), TLR2^-/-^ CTRL (4; n=8).

*Abbreviations*: OF- open field; SD- standard deviation; TST- tail suspension test.

| **Behavioural test: test parameter** | |  |  |  |
| --- | --- | --- | --- | --- |
|  |  | **Mean** | *SD* | *SEM* |
| OF: Duration | 1 | 599.99 | 0.03 | 0.01 |
|  | 2 | 600.00 | 0.00 | 0.00 |
|  | 3 | 600.00 | 0.00 | 0.00 |
|  | 4 | 600.00 | 0.00 | 0.00 |
| OF: Distance | 1 | 28.85 | 5.81 | 1.68 |
|  | 2 | 24.93 | 9.93 | 3.31 |
|  | 3 | 28.11 | 4.67 | 1.21 |
|  | 4 | 29.18 | 10.12 | 3.58 |
| OF: Mean speed | 1 | 0.05 | 0.01 | 0.00 |
|  | 2 | 0.05 | 0.01 | 0.00 |
|  | 3 | 0.05 | 0.01 | 0.00 |
|  | 4 | 0.05 | 0.02 | 0.01 |
| OF: Time mobile | 1 | 486.39 | 43.37 | 12.52 |
|  | 2 | 458.16 | 45.94 | 16.24 |
|  | 3 | 450.40 | 48.80 | 12.60 |
|  | 4 | 421.79 | 64.09 | 22.66 |
| OF: Time immobile | 1 | 113.59 | 43.35 | 12.51 |
|  | 2 | 141.84 | 45.94 | 16.24 |
|  | 3 | 149.60 | 48.80 | 12.60 |
|  | 4 | 178.21 | 64.09 | 22.66 |
| OF: Mobile episodes | 1 | 27.17 | 7.47 | 2.16 |
|  | 2 | 32.25 | 8.36 | 2.96 |
|  | 3 | 35.20 | 8.69 | 2.24 |
|  | 4 | 34.88 | 7.77 | 2.75 |
| OF: Immobile episodes | 1 | 26.67 | 7.67 | 2.21 |
|  | 2 | 32.00 | 8.30 | 2.93 |
|  | 3 | 34.73 | 8.71 | 2.25 |
|  | 4 | 34.13 | 7.68 | 2.72 |
| OF: Line crossings | 1 | 165.58 | 51.73 | 14.93 |
|  | 2 | 139.75 | 38.32 | 13.55 |
|  | 3 | 167.47 | 37.84 | 9.77 |
|  | 4 | 146.38 | 37.18 | 13.14 |
| OF: Absolute turn angle | 1 | 56252.17 | 11455.53 | 3306.93 |
|  | 2 | 43144.38 | 8385.90 | 2964.86 |
|  | 3 | 41261.33 | 8279.64 | 2137.79 |
|  | 4 | 36473.63 | 8784.88 | 3105.92 |
| OF: Max speed | 1 | 0.03 | 0.00 | 0.00 |
|  | 2 | 0.03 | 0.01 | 0.00 |
|  | 3 | 0.03 | 0.00 | 0.00 |
|  | 4 | 0.04 | 0.01 | 0.00 |
| OF: Rotation | 1 | 31.00 | 6.62 | 1.91 |
|  | 2 | 26.38 | 4.44 | 1.57 |
|  | 3 | 34.33 | 7.11 | 1.84 |
|  | 4 | 33.75 | 9.50 | 3.36 |
| OF: Clockwise rotations | 1 | 16.58 | 4.80 | 1.38 |
|  | 2 | 13.25 | 6.69 | 2.37 |
|  | 3 | 16.27 | 6.53 | 1.69 |
|  | 4 | 14.13 | 4.49 | 1.59 |
| OF: Anti-clockwise rotations | 1 | 14.42 | 5.33 | 1.54 |
|  | 2 | 13.13 | 2.75 | 0.97 |
|  | 3 | 18.07 | 7.21 | 1.86 |
|  | 4 | 19.63 | 7.71 | 2.73 |
| YM: Distance | 1 | 48.05 | 9.71 | 2.80 |
|  | 2 | 46.45 | 8.18 | 2.47 |
|  | 3 | 50.62 | 5.75 | 1.66 |
|  | 4 | 42.64 | 4.69 | 1.77 |
| YM: Mean speed | 1 | 0.08 | 0.02 | 0.00 |
|  | 2 | 0.08 | 0.01 | 0.00 |
|  | 3 | 0.08 | 0.01 | 0.00 |
|  | 4 | 0.07 | 0.01 | 0.00 |
| YM: Time mobile | 1 | 440.25 | 42.47 | 12.26 |
|  | 2 | 424.42 | 27.66 | 8.34 |
|  | 3 | 455.93 | 29.42 | 8.49 |
|  | 4 | 413.09 | 13.93 | 5.26 |
| YM: Time immobile | 1 | 159.75 | 42.47 | 12.26 |
|  | 2 | 175.58 | 27.66 | 8.34 |
|  | 3 | 144.08 | 29.42 | 8.49 |
|  | 4 | 186.91 | 13.93 | 5.26 |
| YM: Mobile episodes | 1 | 37.58 | 5.28 | 1.52 |
|  | 2 | 44.36 | 5.75 | 1.73 |
|  | 3 | 34.33 | 4.68 | 1.35 |
|  | 4 | 38.86 | 7.22 | 2.73 |
| YM: Immobile episodes | 1 | 36.50 | 5.32 | 1.53 |
|  | 2 | 44.09 | 5.75 | 1.73 |
|  | 3 | 33.83 | 4.65 | 1.34 |
|  | 4 | 38.29 | 7.13 | 2.70 |
| YM: Line crossings | 1 | 141.08 | 22.47 | 6.49 |
|  | 2 | 140.45 | 28.39 | 8.56 |
|  | 3 | 163.50 | 19.30 | 5.57 |
|  | 4 | 163.14 | 37.62 | 14.22 |
| YM: Absolute turn angle | 1 | 46991.00 | 10471.86 | 3022.97 |
|  | 2 | 43429.45 | 6819.24 | 2056.08 |
|  | 3 | 42262.33 | 7423.61 | 2143.01 |
|  | 4 | 50424.00 | 8257.70 | 3121.12 |
| YM: Max speed | 1 | 0.91 | 0.50 | 0.14 |
|  | 2 | 0.82 | 0.45 | 0.13 |
|  | 3 | 0.57 | 0.06 | 0.02 |
|  | 4 | 0.53 | 0.08 | 0.03 |
| YM: Rotations | 1 | 20.42 | 4.06 | 1.17 |
|  | 2 | 21.64 | 4.65 | 1.40 |
|  | 3 | 26.75 | 3.17 | 0.91 |
|  | 4 | 25.57 | 2.07 | 0.78 |
| YM: Clockwise rotations | 1 | 12.83 | 5.04 | 1.46 |
|  | 2 | 12.64 | 5.39 | 1.63 |
|  | 3 | 15.33 | 3.98 | 1.15 |
|  | 4 | 11.00 | 3.87 | 1.46 |
| YM: Anti-clockwise rotations | 1 | 7.58 | 3.78 | 1.09 |
|  | 2 | 9.00 | 5.04 | 1.52 |
|  | 3 | 11.42 | 4.80 | 1.38 |
|  | 4 | 14.57 | 4.43 | 1.67 |
| YM: Path efficiency (%) | 1 | 1.05 | 0.58 | 0.16 |
|  | 2 | 1.09 | 0.66 | 0.20 |
|  | 3 | 0.625 | 0.49 | 0.14 |
|  | 4 | 1.04 | 0.53 | 0.20 |
| YM: Spontaneous alternation % | 1 | 63.24 | 5.65 | 1.63 |
|  | 2 | 61.03 | 9.48 | 2.86 |
|  | 3 | 59.65 | 6.02 | 1.74 |
|  | 4 | 58.55 | 4.36 | 1.65 |
| TST: Duration | 1 | 360.00 | 0.00 | 0.00 |
|  | 2 | 360.00 | 0.00 | 0.00 |
|  | 3 | 360.00 | 0.00 | 0.00 |
|  | 4 | 358.81 | 3.14 | 1.19 |
| TST: number of presses | 1 | 33.86 | 11.94 | 3.19 |
|  | 2 | 31.33 | 10.17 | 3.39 |
|  | 3 | 28.33 | 9.56 | 2.76 |
|  | 4 | 36.00 | 8.16 | 3.09 |
| TST: Mirovanje : time pressed | 1 | 103.72 | 48.09 | 12.85 |
|  | 2 | 152.68 | 34.69 | 11.56 |
|  | 3 | 99.43 | 33.17 | 9.57 |
|  | 4 | 144.90 | 56.08 | 21.20 |
| TST: latency 1st press | 1 | 27.65 | 7.11 | 1.90 |
|  | 2 | 27.37 | 12.79 | 4.26 |
|  | 3 | 45.04 | 22.08 | 6.37 |
|  | 4 | 30.14 | 12.38 | 4.68 |
| TST: latency 1st release | 1 | 28.15 | 7.44 | 1.99 |
|  | 2 | 27.81 | 13.16 | 4.39 |
|  | 3 | 45.42 | 22.18 | 6.40 |
|  | 4 | 30.99 | 11.84 | 4.48 |
| TST: longest press | 1 | 18.28 | 10.79 | 2.88 |
|  | 2 | 23.10 | 7.84 | 2.61 |
|  | 3 | 17.87 | 6.89 | 1.99 |
|  | 4 | 17.80 | 6.76 | 2.55 |
| TST: shortest press | 1 | 0.11 | 0.09 | 0.02 |
|  | 2 | 0.12 | 0.08 | 0.03 |
|  | 3 | 0.11 | 0.07 | 0.02 |
|  | 4 | 0.10 | 0.06 | 0.02 |
| TST: mean press duration | 1 | 3.30 | 1.82 | 0.49 |
|  | 2 | 5.16 | 1.53 | 0.51 |
|  | 3 | 3.53 | 1.13 | 0.33 |
|  | 4 | 4.10 | 1.79 | 0.68 |
| TST: press frequency | 1 | 0.09 | 0.03 | 0.01 |
|  | 2 | 0.09 | 0.03 | 0.01 |
|  | 3 | 0.08 | 0.03 | 0.01 |
|  | 4 | 0.10 | 0.02 | 0.01 |

### 6.Weight Changes

##### Table S7. Timeline of changes in weight is shown in four investigated groups. Values are shown as percentage of change from the baseline weight ± standard deviation. n denotes number of animals per group; animals were grouped according to their genotype and intervention. Groups: 1 = TLR2 IH; 2= TLR2 CTRL; 3= TLR2^-/-^ IH; 4= TLR2^-/-^ CTRL; Abbreviations: SD- standard deviation.

| **∆ Weight (%)** | | **Mean** | **SD** |
| --- | --- | --- | --- |
| Third day *vs.* baseline | 1 | -6.27 | 5.39 |
|  | 2 | 1.17 | 2.86 |
|  | 3 | -4.75 | 2.32 |
|  | 4 | -1.50 | 1.20 |
| Sixth day *vs* baseline | 1 | -4.53 | 4.94 |
|  | 2 | 3.58 | 2.39 |
|  | 3 | -3.31 | 2.36 |
|  | 4 | 0.63 | 1.41 |
| Ninth day *vs* baseline | 1 | -4.27 | 5.13 |
|  | 2 | 6.25 | 10.14 |
|  | 3 | -3.81 | 2.17 |
|  | 4 | 0.00 | 1.31 |
| Weight difference 16^th^ day *vs.* baseline | 1 | -3.33 | 6.20 |
|  | 2 | 7.64 | 4.54 |
|  | 3 | -4.33 | 2.77 |
|  | 4 | 2.88 | 4.61 |
| Weight difference 21^st^ day *vs.* baseline | 1 | -2.00 | 6.19 |
|  | 2 | 5.60 | 4.34 |
|  | 3 | -4.89 | 3.76 |
|  | 4 | 0.33 | 1.03 |

##### Table S8. Significance of differences in weight changes during exposure to intermittent hypoxia versus those in their respective controls is shown for groups with functional (TLR2) and non-functional gene (TLR2^-/-^). Post-hoc Mann- Whitney U-test calculated P values of data presented in S7.

| **∆ Weight** | **TLR2 (*P*)** | **TLR2^-/-^ (*P*)** |
| --- | --- | --- |
| Third day vs. baseline | 0.001 | 0.023 |
| Sixth day vs baseline | <0,001 | 0.007 |
| Ninth day vs. baseline | <0,001 | 0.009 |
| 16^th^ day vs. baseline | <0,001 | 0.042 |
| 21^st^ day vs. baseline | 0.033 | 0.031 |

***References:***

11.	Wood TC QUIT: QUantitative Imaging Tools.

20.	Hanish AE, Butman JA, Thomas F, Yao J, Han JC (2015): Pineal hypoplasia, reduced melatonin and sleep disturbance in patients with PAX6 haploinsufficiency. *Journal of sleep research*.

79.	Liu A, Jain N, Vyas A, Lim LW (2015): Ventromedial prefrontal cortex stimulation enhances memory and hippocampal neurogenesis in the middle-aged rats. *eLife*. 4.

80.	Mehedint MG, Craciunescu CN, Zeisel SH (2010): Maternal dietary choline deficiency alters angiogenesis in fetal mouse hippocampus. *Proceedings of the National Academy of Sciences of the United States of America*. 107:12834-12839.

81.	Lim DC, Brady DC, Soans R, Kim EY, Valverde L, Keenan BT, et al. (2016): Different Cyclical Intermittent Hypoxia Severities have Different Effects on Hippocampal Microvasculature. *Journal of applied physiology*.jap 01040 02015.

82.	Squillario M, Barla A (2011): A computational procedure for functional characterization of potential marker genes from molecular data: Alzheimer's as a case study. *BMC medical genomics*. 4:55.
